# Characterizing the microbial metagenome of calcareous stromatolite formations in the San Felipe Creek in Anza Borrego Desert

**DOI:** 10.1101/2023.05.12.540589

**Authors:** Rosalina Stancheva, Arun Sethuraman, Hossein Khadivar, Jenna Archambeau, Ella Caughran, Ashley Chang, Brad Hunter, Christian Ihenyen, Marvin Onwukwe, Dariana Palacios, Chloe La Prairie, Nicole Read, Julianna Tsang, Brianna Vega, Cristina Velasquez, Xiaoyu Zhang, Elinne Becket, Betsy Read

## Abstract

Here we describe the metagenome composition, community functional annotation, and diversity of prokaryotic microbial species derived from calcareous stromatolite formations discovered in the dry stream bed of the open-canopy, ephemeral San Felipe Creek in the Anza Borrego Desert. In this environment, resident microbes must be able to adapt to the harsh conditions of extreme heat, high UV light, desiccation and fluctuating solubilization/precipitation and hydration/evaporation. Metagenomic analysis revealed a community capable of carrying out complete nitrogen fixation and assimilatory nitrate reduction, forming biofilms and quorum sensing, and potentially forming thick-walled akinetes as desiccation-resistant stages. Nitrogen cycling is likely to play a fundamental role in mediating both the structure of the stromatolite microbial community and the mineral precipitation/dissolution. The viruses present in the stromatolites, particularly *Nodularia* and *Mycobacterium* phages are also likely to impact community population dynamics and activity. Stromatolite community members possess different morphological and physiological strategies to cope with desiccation stress.

Metagenomic signatures were found for scytonemin, carotenoids, synthesis of potential microsporine-like amino acids; genes involved in microalgal desiccation tolerance, including those encoding aquaporins, chaperones, antioxidants; and enzymes responsible for the synthesis of trehalose, sucrose, and polyamines.

The stromatolite ecosystem provides a diverse array of microniches where different functional guilds can develop complex metabolite exchange with the substrate supporting their life in extreme conditions. Metagenome analyses revealed several genes that might enable a specialized and unique group of endolithic cyanobacteria including *Chroococcidiopsis, Hyella, Myxosarcina*, and *Pleurocapsa* to derive metals and important nutrients from rocks, being potentially destructive for the calcareous formations. Our study revealed environmental adaptations of freshwater microbial communities in desert stream stromatolites which may provide valuable insights into Precambrian paleoenvironments, which are little known.

## Introduction

Stromatolites are laminated or lithified organosedimentary structures produced over an extended period from the activity of microbes including photosynthetic cyanobacteria. The formation of the unique structures occurs through a process that begins with photosynthetic carbon assimilation, the development of biofilms rich in extracellular polysaccharides, and initial supersaturation of the macroenviroment with CaCO3 minerals. The biofilms serve as precursors to seed crystal formation and facilitate the trapping and binding of sediments during the mineralization process that result in the formation of calcareous stromatolites (Arp et al. 2001). As complex self-sustained ecosystems that develop on solid/water interfaces, stromatolites represent some of the earliest forms of marine microbial life on Earth, dating back some 3.5 billion years (Schopf, 2006; Schopf and Packer, 1987). The first microbial stromatolite ecosystems most likely formed under hypersaline or intertidal marine conditions as indicated by geological evidence for higher salinity of the Precambrian (> 550 millions year ago) ocean (Arp et al. 2001). One of the oldest non-marine carbonate stromatolites formed by fluvial-lacustrine microbial communities were recorded within the Mesoproterozoic (1.09 billion years ago) Copper Harbor Conglomerate of northern Michigan (Elmore, 1983). However, little is known about the Precambrian freshwater paleoenvironments and how microbial communities forming stromatolites were adapted to various environmental conditions, such as UV radiation, desiccation stress, changes in alkalinity, and variable discharge (Fedorchuk et al. 2016).

Living stromatolites are rare compared to their fossilized counterparts and are recorded in select geographically isolated locations in both marine and freshwater environments. Some of the most well-known living stromatolites are distributed in the hypersaline coastal mats of the Hamelin Pools of Shark Bay in Western Australia (Playford and Cockbain, 1976); in the open ocean Exuma Cays in the Bahamas (Reid et al., 1995); in the nonhalophilic nonthermophilic oligotrophic lakes in the Cuatro Cienagas Basin in Chihuahuan Desert, Coahuila, Mexico (Falcon et al., 2007), in fluvio-lacustrine systems in Ruidera Pools National Park in Central Spain (Santos et al. 2010), and in the hot springs in Yellowstone National Park (Walter et al.,1972; Berelson et al., 2011). Many recent stromatolite freshwater habitats are extreme and periodically exposed to desiccation events (Santos et al. 2010), and as such are considered environmental analogues to those found in Precambrian shallow waters (Foster and Mobberley, 2010).

While marine and freshwater stromatolites are similar in terms of their overall structure and the manner in which they are formed, the microbial community composition is expected to differ. The microbial community in stromatolites plays an important role in the biogeochemical cycling of nutrients and in the maintenance and structure of these unusual ecosystems. Their composition and diversity vary and reflects the prevailing environmental conditions under which they are formed. In freshwater ecosystems, stromatolites are typically found in shallow, placid bodies of water such as lakes, ponds, and streams where factors such as minerals, nutrients, light, and water dynamics can be extreme and impact the microbial inhabitants (Proemse et al., 2017; Santos et al., 2010; Farias et al., 2013; Wacey et al., 2018). For example, in the Quatro Cienegas Basin, oligotrophy and habitat disturbance appear to be important factors governing the formation and community structure of the stromatolites. The metagenomic signatures of microbial mat forming stromatolites in permanent green pools in Quatro Cienegas Basin, differ significantly from those in the red desiccation ponds, despite residing in similar oligotrophic bodies of waters with limiting phosphate. The stromatolite microbial community in the green pools tends to be more diverse and dominated by phototrophic Cyanobacteria (orders Chroococcales and Nostocales). Whereas heterotrophic members of the metabolically opportunistic Proteobacteria and Bacteroidetes dominate the microbial mats in the red ponds where the water levels are lower and higher alkalinity and elevated solute concentrations are expected (Bonilla-Rosso et al., 2012). The microbial communities of marine stromatolites found in shallow coastal regions, coral reefs, or deep-sea hydrothermal vents are dependent upon environmental conditions such as salinity, temperature, and nutrients. These formations are also dominated by bacteria and are equally as diverse and complex as their freshwater counterparts. For instance, the most prominent phyla present in stromatolites found across different locations in Shark Bay include Proteobacteria (55-69%), Cyanobacteria (15-29%), Plantomycetes (5-7%), and Bacteroidetes (3-7%), Verrucomicrobia (2-3%) Chloroflexi (2-5%), and Actinobacteria (0.25-2%) (Babilonia et al., 2018).

Our study area, Southern California has a Mediterranean climate with cool, wet winters and hot, dry summers (Luo et al. 2017) which determines the intermittent flow regime of many local streams. The biotic communities of non-perennial streams, which cease to flow at some point in time or space, are largely defined by drying disturbances (Busch et al. 2020) and contain many benthic algae which are adapted to desiccation (Stancheva et al. 2019). These streams have alkaline low-nutrient water and two distinct states: wet with flowing surface water, and dry (without flow) state with duration from nearly month to a year in southern California (Busch et al. in press). Mediterranean calcareous streams are usually highly productive (Sabater et al. 2017). For instance, the median species richness of soft-bodies algal communities in non-perennial streams in southern California (25 species) was two-fold higher than the median algal species richness (12 species) recorded in 104 perennial and non-perennial streams in the same area (Stancheva et al. 2012, Busch et al. in press).

Stromatolite-like carbonated, characteristically laminated formations on the streambed were described from Mediterranean calcareous streams in Europe. The stromatolite communities were shaped by cyanobacteria (e.g. *Rivularia* and *Schizothrix*) with some contributions from green algae and diatoms, formed in open-canopy streams with intense photosynthetic production and associated calcium carbonate precipitation (Sabater et al. 2000).

Hot deserts, such as Chihuahuan Desert in northern Mexico, and Anza Borrego Desert in southern California offer limited freshwater habitats to organisms adapted to survive the extreme variability in precipitation and other restrictive environmental factors. Most of the streams in these deserts are ephemeral and their flow is tightly linked to short rainfall episodes. We studied an open-canopy desert San Felipe Creek in the Anza Borrego Desert, Southern California, which is completely dry for long periods of time. Large areas of the dry stream bed of the creek are covered by persistent stromatolite-like laminated calcareous structures. Living stromatolites were previously reported in ephemeral sections of alkaline streams in the Anza Borrego Desert (Buchham 1995) where there is maximum evaporation and high solute concentrations. Here we characterize the microbial metagenome of laminated stromatolite formations in the Anza Borrego Desert, to address (1) the composition of the microbial community in the Anza Borrego Desert stromatolites, (2) the functional roles of the resident microbes, and 3) how they interact.

## Materials and methods

Materials were collected on November 20, 2019 from top of granite rocks in the ephemeral San Felipe Creek (Figure 1) (coordinates: 33.0986, -116.4708) location nearby previously studied locality of living stromatolites (Buchheim et al., 1993). Dry stromatolites (2 cm thick) were chiseled from the surface of the rock with gloved hands and placed in sterile 50 mL falcon tubes. Samples were transported to California State University San Marcos where they were stored at room temperature. San Felipe Creek was dry at the time of collecting the stromatolite samples, therefore we conducted additional field trips until water was observed in small pools on November 13, 2022 and January 26, 2023 allowing water chemistry measurements to be obtained (temperature, conductivity, salinity, total dissolved solids) with anExSick II ES400 field meter (Nashua, NH, USA). We used temperature and precipitation data for the closest NOAA Station Borrego Desert Park (coordinates 33.2559°, -116.4036°) from National Weather Service https://www.weather.gov/wrh/climate?wfo=sgx.

**Figure 1.**
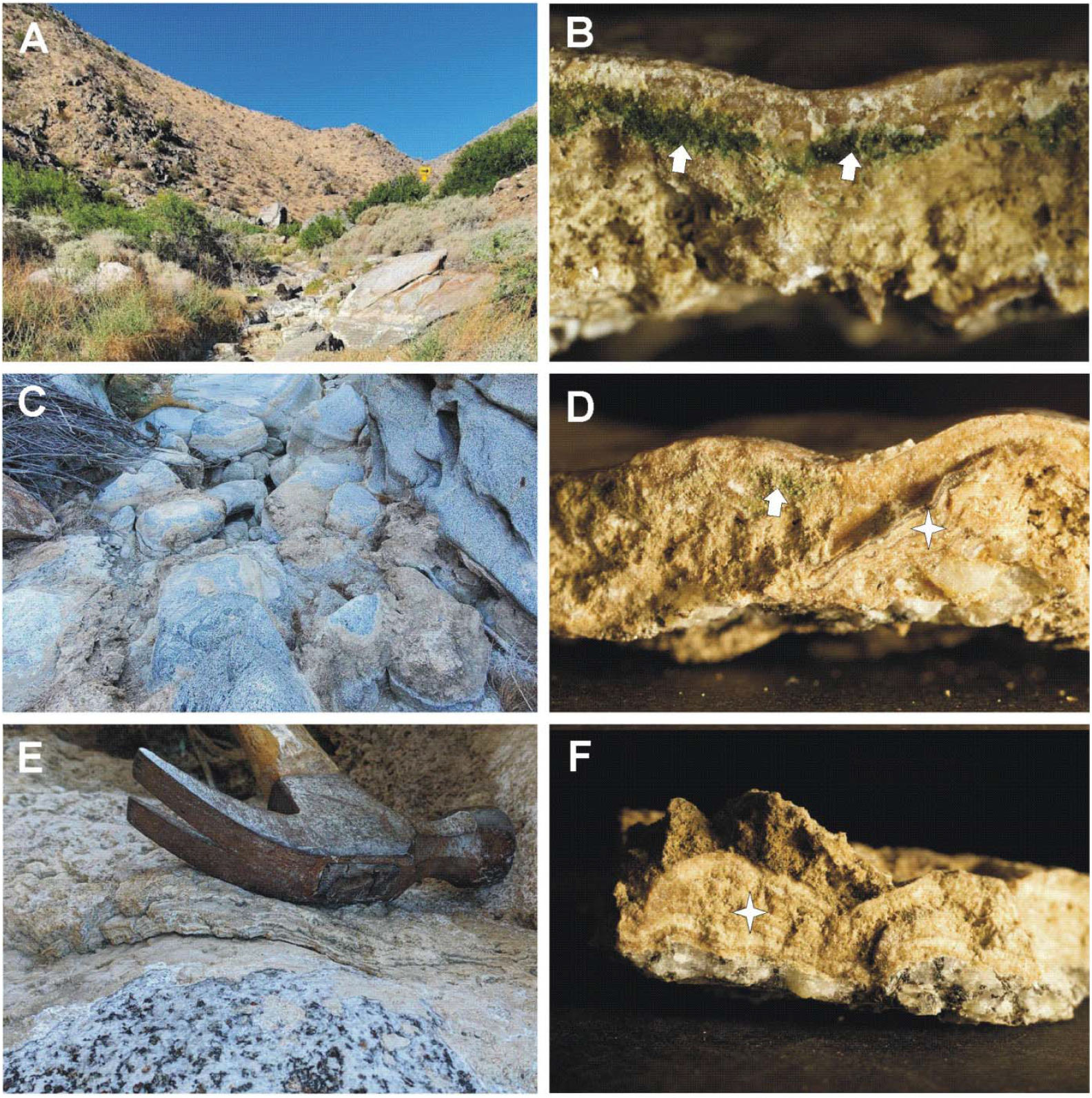
The ephemeral San Felipe Creek in Anza Borrego Desert **(A)**. Samples were collected from the surface of boulders found in bed of the desiccated creek **(C, E)**. Samples imaged under a dissecting scope **(B, D, F)** show characteristic features of stromatolites including endolithic cyanobacteria underneath a layer of calcite (arrows in B and D) and multiple laminate layers of organic and inorganic material (asterisks in D and F). DNA was isolated from the stromaolites shown in panels B and E.

Dry stromatolite material was crushed with sterile ceramic beads using a Fisherbrand Bead Mill 24 Homogenizer (Fisher Scientific) with a four-cycle agitation protocol (one minute 26 “on”, one minute “off”) at 3000 rpm. Thereafter, using 500 mg of ground material and following the protocol of Narayan et al., 2016, cells were extracted with a PEG-NaCl buffer. Cellular lysis was accomplished by applying a combination of enzymatic and hot detergent incubations.

Metagenomic DNA was then isolated by selective precipitation. The concentration of DNA was determined by absorbance at A260, while the purity was verified by the A260/280 and A260/230 ratios on a nanodrop spectrophotometer and the integrity confirmed by agarose gel electrophoresis. Four replicate metagenomic libraries were prepared using the Transposase Enzyme Linked Long-read Sequencing (TELL-Seq) Whole Genome Sequencing (WGS) Library Prep Kit from Universal Sequencing (Carlsbad, CA). Briefly, approximately 5 ng of input DNA was barcoded using 16 million TELL beads per sample. After capture and clean up, the metagenomic libraries were PCR amplified using 12 cycles. The libraries were then pooled and sequenced on the Illumina(R) NextSeq 500/550 platform using the Mid Output Kit V2.5 (150 cycles).

Raw reads were compiled as FASTQ files, and quality of sequencing was assessed using FastQC v.11.9 (ref) and a conservative PHRED Q score cutoff of 30 was used to trim reads during the de novo metagenome assembly process using the UST TELL-Seq assembly pipeline. QUAST v.4.4 (Gurevich *et al*. 2013) was used to assess quality of the metagenome assemblies using default parameters on KBase v.2.6.4 (Allen *et al*. 2017), www.kbase.us).

Taxonomic classification was performed at four different levels: (1) raw sequencing reads using Kaiju v.1.7.3 (Menzel *et al*. 2016 against the NCBI microbial genomes database in Kbase v.2.6.4 (Allen *et al*. 2017), (2) metagenome assemblies using Kraken2 v.2.0.8 (Wood *et al*. 2019) with the GTDB database release 202 (Parks et al., 2018), (3) contig binning using CONCOCT (Alneberg et al., 2014) and MaxBin2 (Wu et al., 2016), and (4) protein coding genes using GhostKOALA v.2.2 (Kanehisa et al., 2016) against the Kyoto Encyclopedia of Genes and Genomes (KEGG) database. Raw reads from all metagenome libraries were normalized using scaling with ranked subsampling (Beule and Karlovsky 2020). To assess the diversity and relative composition of the assembled metagenomes, alpha indices (Shannon, Simpson, Chao) were computed using the normalized mapped read counts in R v.4.2.1. Automated gene calling and functional analyses were performed using the NCBI Prokaryote Genome Annotation Pipeline (PGAP; Tatusova et al., 2016) with *Lyngbya aestuarii* and *Sediminibacterium* as references. All PGAP run parameters are specified in the scripts accessible on the project’s GitHub page. Predicted proteins were classified using GhostKOALA v.2.2 (Kanehisa et al., 2016) to obtain functional capabilities of the microbial communities and to assess interactions between identified organisms.

## Results and Discussion

San Felipe Creek is an open-canopy ephemeral stream in the Anza Borrego Desert, which is dry most of the year. California experienced a drought in 2011-2017, altering vegetation cover in the desert creeks. Mean annual total precipitation in Borrego Park Station for the study period (2019-2023) was 4.99 in (range from 2.76 in for 2022 to 9.25 in for 2019). Mean monthly lowest min temperature was 36° F (range 34° F in January 2019 to 77° F in August 2022). Mean monthly highest max temperature was 113° F (range 72° F in January 2023 to 121° F in July 2020).

During the course of this study, we visited the stream several times between November 2019 and January 2023. The stream received some water in the winter rain season that was retained for very short periods of time. The water chemistry parameters measured on November 13, 2022 and January 26, 2023 in a very small water pool remaining after a winter rainstorm were water temperature 11.8 - 13.8° C, pH 8.35, conductivity 6830 - 7310 µS/cm2, salinity 3.77 - 4 ppt, total dissolved solids 4.7 - 5.13 ppt. Our measurements showed elevated ion content and salinity within the brackish water range. Buchheim (1993) studied stromatolites in two creeks in Anza Borrego Desert (e.g. San Felipe Creek and nearby Carrizo Creek) following major flood in the spring of 1993. In the months following the flood he recorded in Carrizo Creek a water depth of 0.1 to 1.5 m, pH 8.4-8.5, Chlorides 209-396 ppm, Na 207-281 ppm, Ca 106- 169 ppm, SO_4_ 218-682 ppm, which were three times higher in previous visit in May 1990, illustrating the importance of water availability on water chemistry and biota in these ephemeral streams.

### Metagenome Sequencing and Assembly

Pooled sequencing of all four libraries generated an average of 2.05 × 10^7^ reads with an average GC content of 54.5% (Supplementary Table 1), with over 90.6% of reads exceeding an average PHRED quality score of 30. After trimming for Q > 30, individual libraries were assembled to produce separate metagenome assemblies (Supplementary Table 1). Despite the approximately 2.8 Gb of sequence data generated from each of the replicate libraries, the average metagenome assembly was just 1.3 Mb, with a sequence depth of 10X. This coupled with the small number of scaffolds >1000 bp (x = 145), the comparatively large L50 (x = 654), and small N50 (x= 652) reflects highly fragmented and incomplete metagenomes (Supplementary information Table 1). When reads from the replicate libraries were combined and the assembly were optimized using K-mers of 47, a larger more contiguous metagenome was yielded at ∼29 Mb. The largest scaffold was ∼1.1 Mb with an L50 count of 109 and an N50 size of 56 Kb (Table 1).

**Table 1.**
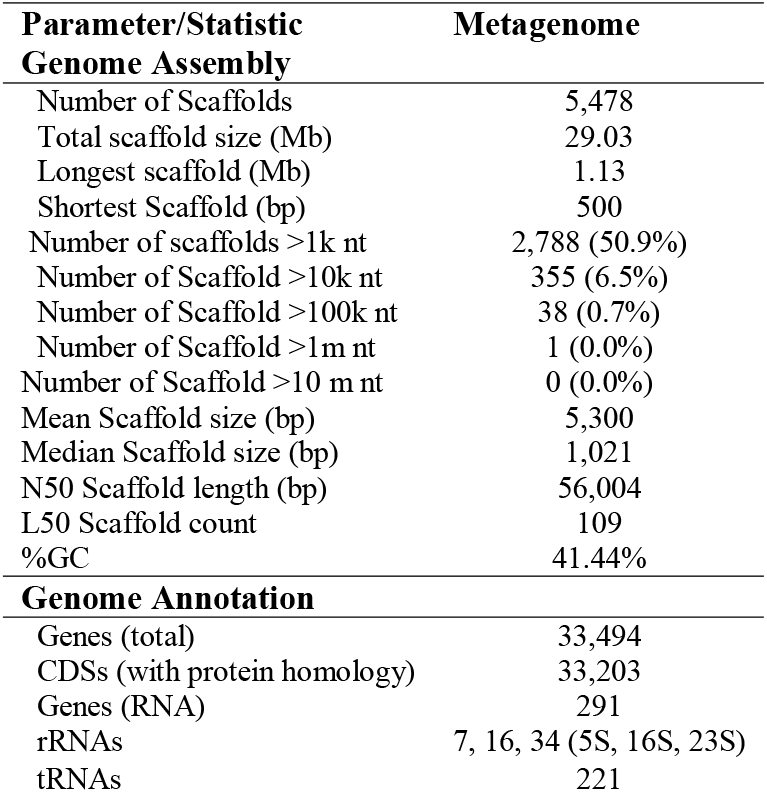
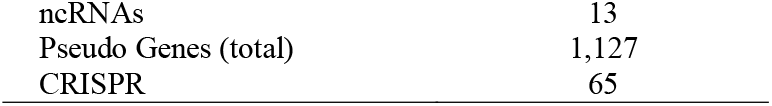
Quality metrics were performed on the combined metagenomics data using QUAST and show a relatively large and contiguous 29 Mb metagenome.

To determine the microbial composition and putative ecological function of members of the stromatolite community, unsupervised contig binning of the metagenome was performed. When binning was accomplished by coverage and composition using CONCOCT (Alneberg et al., 2014), 84 bins were returned that covered ∼93% of the metagenome, however, all were deemed “inconclusive” and only eight yielded identities >80% with the query coverages greater than 50% (Supplementary Table 2). Although deemed “inconclusive”. All the assigned metagenome-assembled genomes (MAG) were predicted to be of cyanobacterial origin.

Seven MAGs identified belonged to the order *Nostacales* and are estimated to be either *Nostoc punctiforme* or *Tolypothrix tenuis*, and one affiliated with *Oscallatoriophycideae* projected to be *Rubdibacter lacunae* (*R. lacunae*). *R. lacunae* is a phycoerythrin-containing cyanobacteria isolated from seawater, which is interesting given the hypersaline conditions of the San Felipe Creek. MaxBin2 which uses composition together with abundance and phylogenetic markers (Wu et al., 2016) to bin contigs yielded 5 bins that covered just 15% metagenome. All of the MAGs were all regarded as “inconclusive”, with predicted organisms returned for only three of the bins, all of which were *Nostoc punctiforme*. The poor results from both CONCOCT and MaxBin2 suggest that sequence coverage is insufficient and/or that the microdiversity of the sample is high. While the sequence reflects an incomplete metagenome and the community and metabolic activities can only be partially described, the information presented is the first of its kind for a unique stromatolite site that exists in a freshwater spring in the Anza Borrego desert.

### Community Composition

Classification of OTUs on the raw reads identified communities were primarily comprised of Cyanobacteria (85.95%), followed by Proteobacteria (5.73%) and Actinobacteria (4.33%) (Figure 2). Analyses of community composition based on the metagenome assemblies showed a similar trend to that of the raw reads, with the vast majority of contigs mapped to Cyanobacteria and remaining phyla less than two percent abundant (Figure 2A). Genus level classification had a higher abundance of *Nostoc* and *Aulosira* when the raw reads were assembled (Figure 2B). The increase of major phylum and genera from raw reads to assemblies is likely a result of the high abundance of cyanobacteria lowering the relative abundances of other key players. Cyanobacteria account for 98.29% of the assembly, which has potentially masked other phyla. This is evident in the genera present above two percent abundance, all of which are part of the Cyanobacteria phylum.

**Figure 2.**
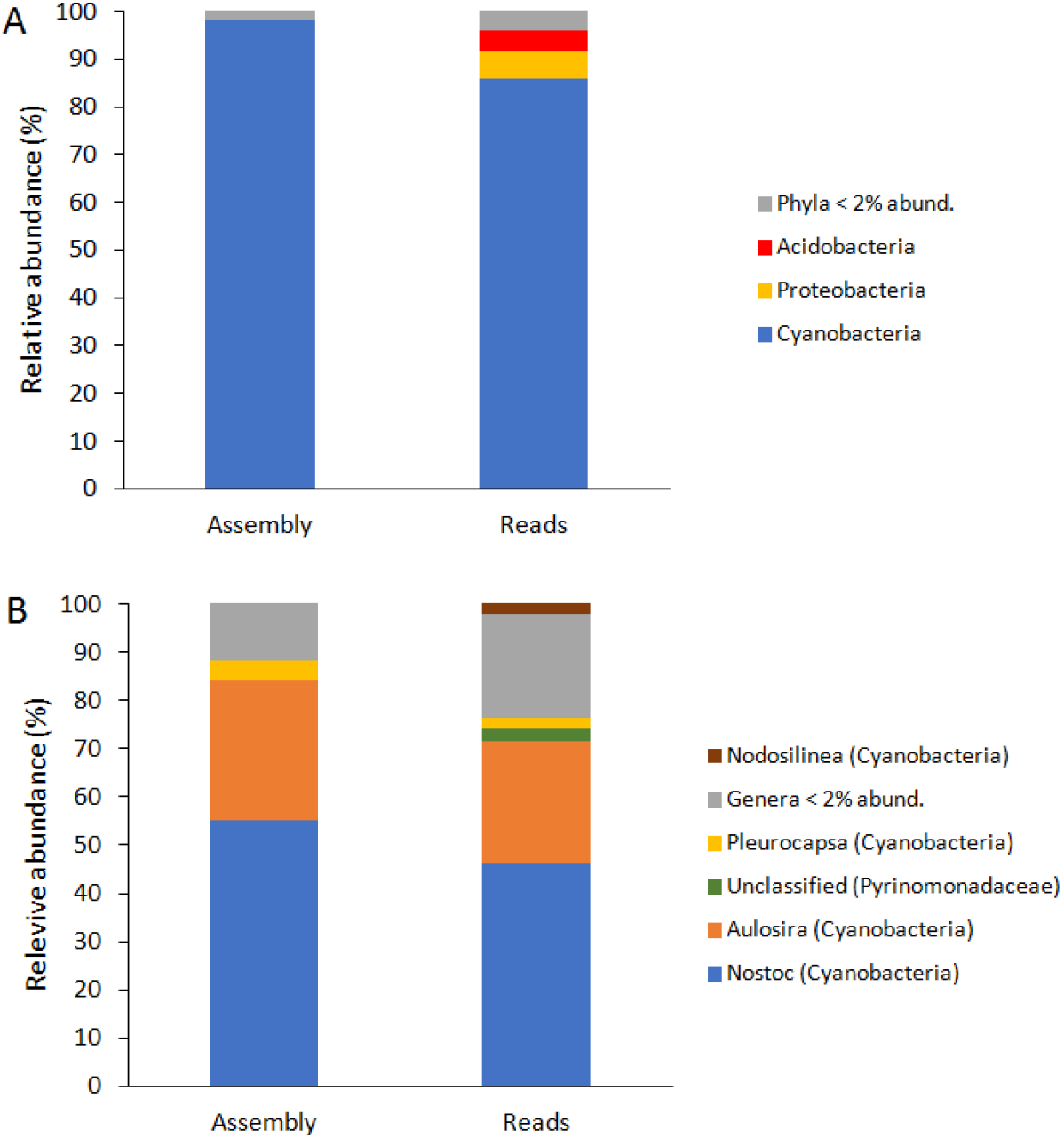
Taxonomic composition of the microbial community in the stromatolites of the San Felipe Creek in Anza Borrego is dominated by cyanobacteria at phylum (A) and genus (B) level. Data for the complete prokaryote microbial community for the assembled metagenome and raw sequence reads are presented.

We identified 45 cyanobacterial genera, belonging to 7 orders and 6 unclassified cyanobacteria which make up 95.55% of total cyanobacterial abundance (Figure 3, see Table 2 for complete list of genera). They belong to orders Nostocales (19 genera, 91.84% of total gene copies), Synechococcales (8 genera, 0.81%), Oscillatoriales (5 genera, 0.21%), Pleurocapsales (6 genera, 4.82%), Chroococcidiopsidales (3 genera, 0.26%), Chroococcales (3 genera, 0.26%), and Gloeomargaritales (1 genus, 0.05%). Within the stromatolite metagenome, both heterocytous (order Nostocales) and non-heterocytous cyanobacteria are present. The most abundant genera were *Nostoc* (55.29%), *Aulosira* (29.16%), *Pleurocapsa* (3.93%) and *Trichormus* (1.83%). The rest of the cyanobacterial genera were below 2% abundance. The cyanobacterial community in the stromatolites was dominated by heterocytous species belonging to order Nostocales (91.78%), presumably able to fix atmospheric nitrogen. This cyanobacterial community is indicative for nitrogen limiting conditions previously documented in stream in southern California where the nitrogenase gene expression in similar cyanobacteria taxa (e.g. *Nostoc* and *Calothrix*) was demonstrated by real time PCR (Stancheva et al. 2013). Certain types of cyanobacteria in the stromatolites such as *Calothrix* and *Rivularia*, possess heteropolar filaments that end in lengthy, colorless strands that serve as sites for external phosphatase activity. In environments where phosphorus is scarce, cyanobacteria benefit from having long hairs that enable them to forage for organic phosphorus when inorganic phosphorus is depleted. Stancheva et al. (2013, Fig. 2C) showed that *Rivularia* reached its highest abundance in stream sites with high N:P ratio in contrast to the rest of N_2_-fixing cyanobacteria which flourish in N:P<15 thus nitrogen-limited conditions.

**Figure 3.**
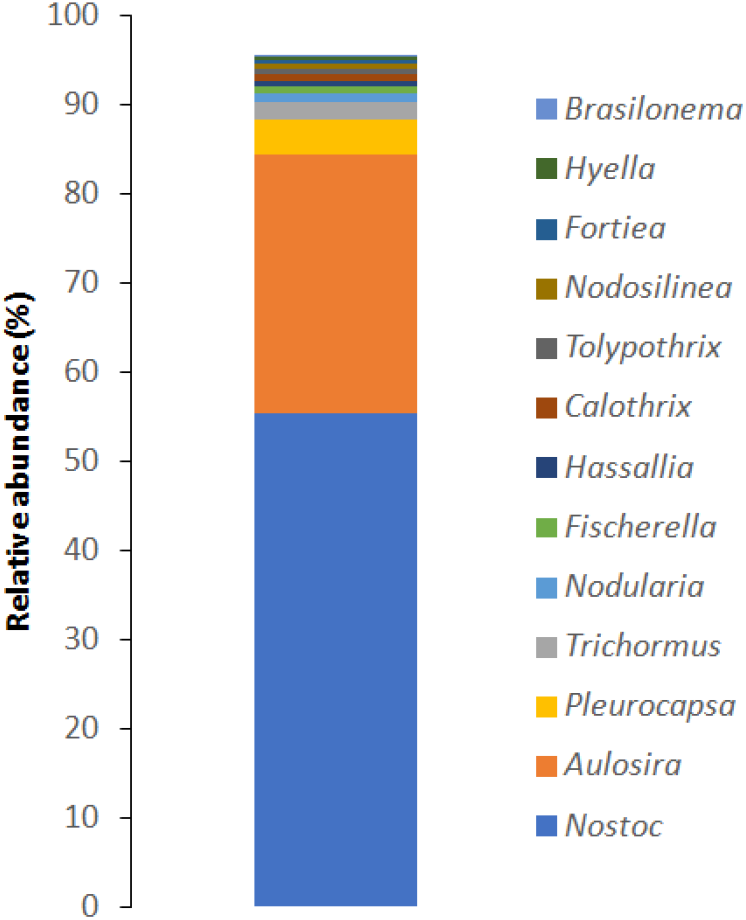
Top 13 most abundant genera of Cyanobacteria in the assembled genome of the stromatolites from San Felipe Creek in the Anza Borrego Desert.

**Table 2.**
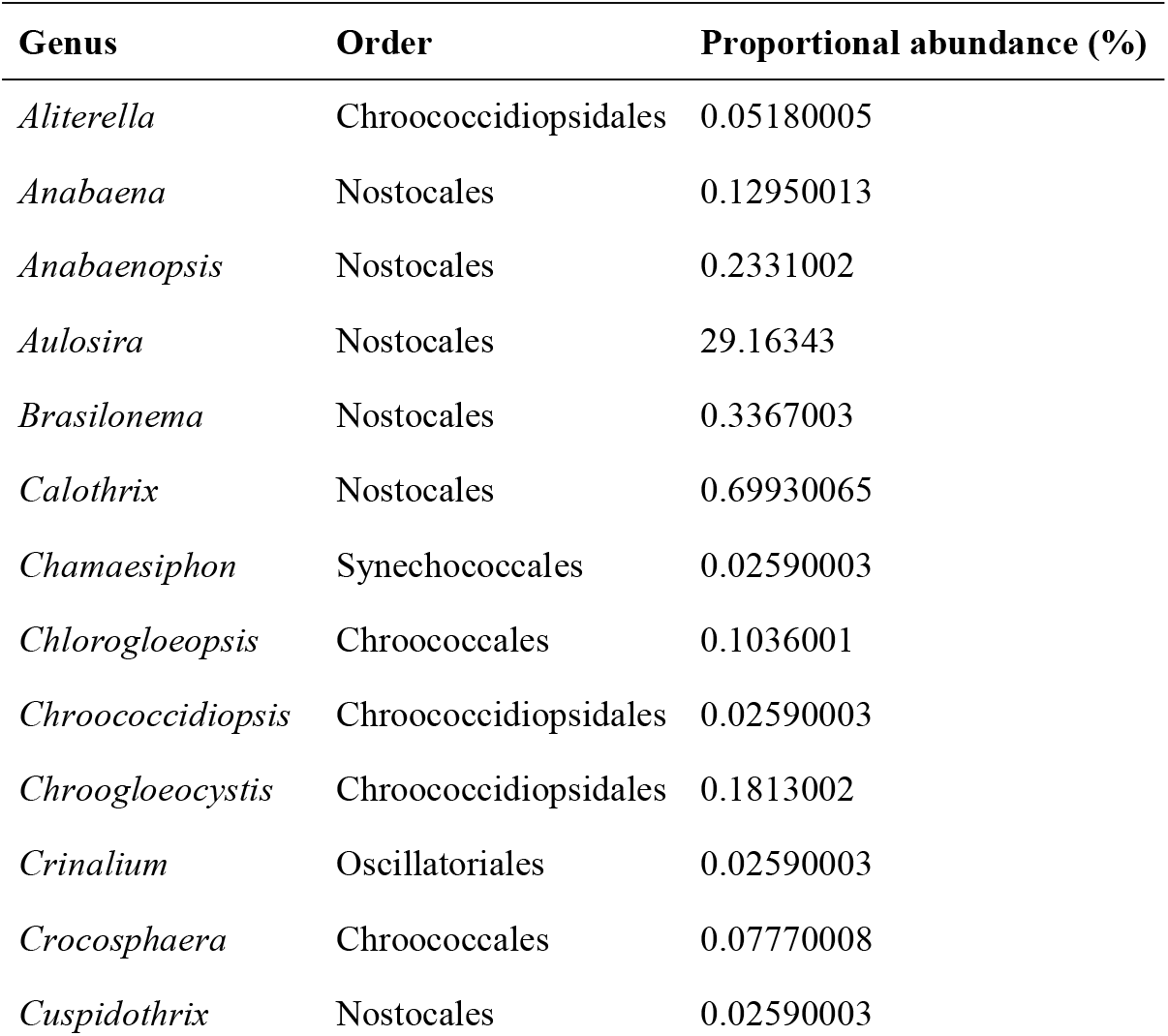
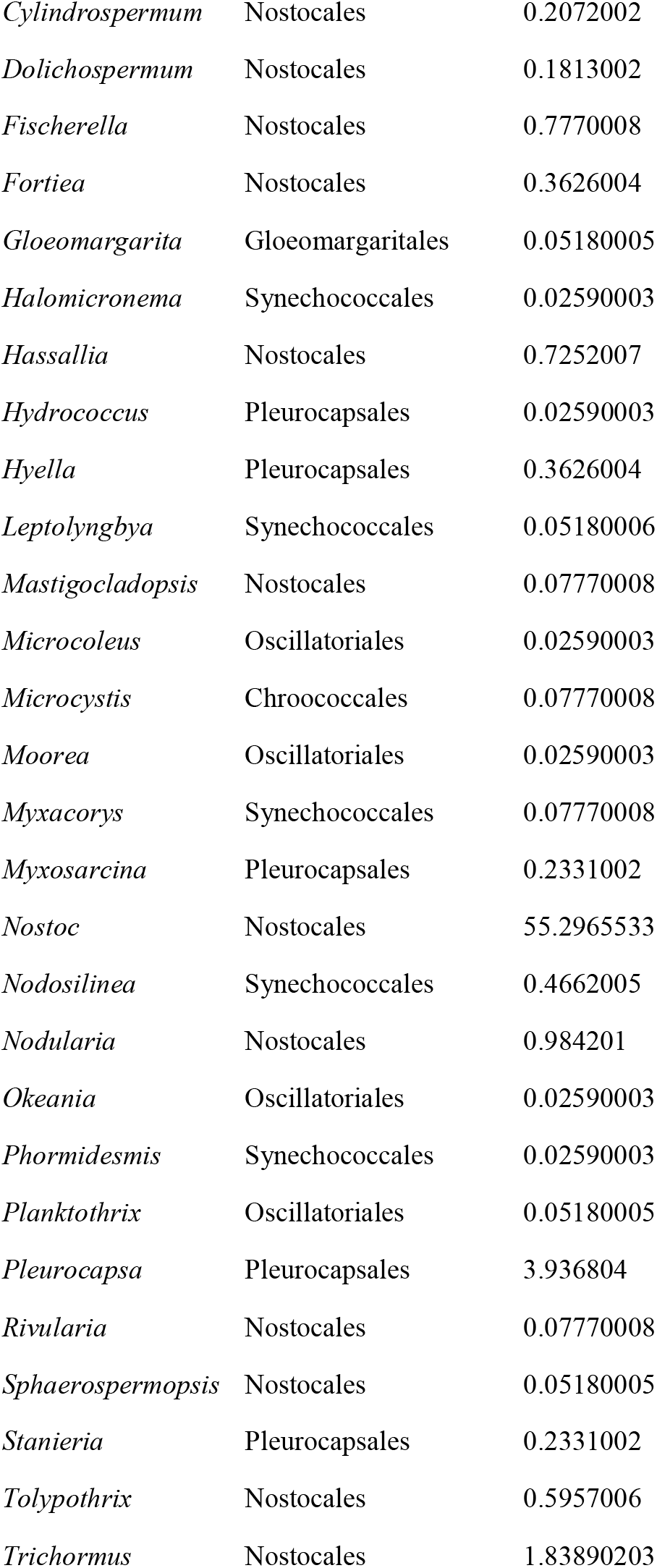
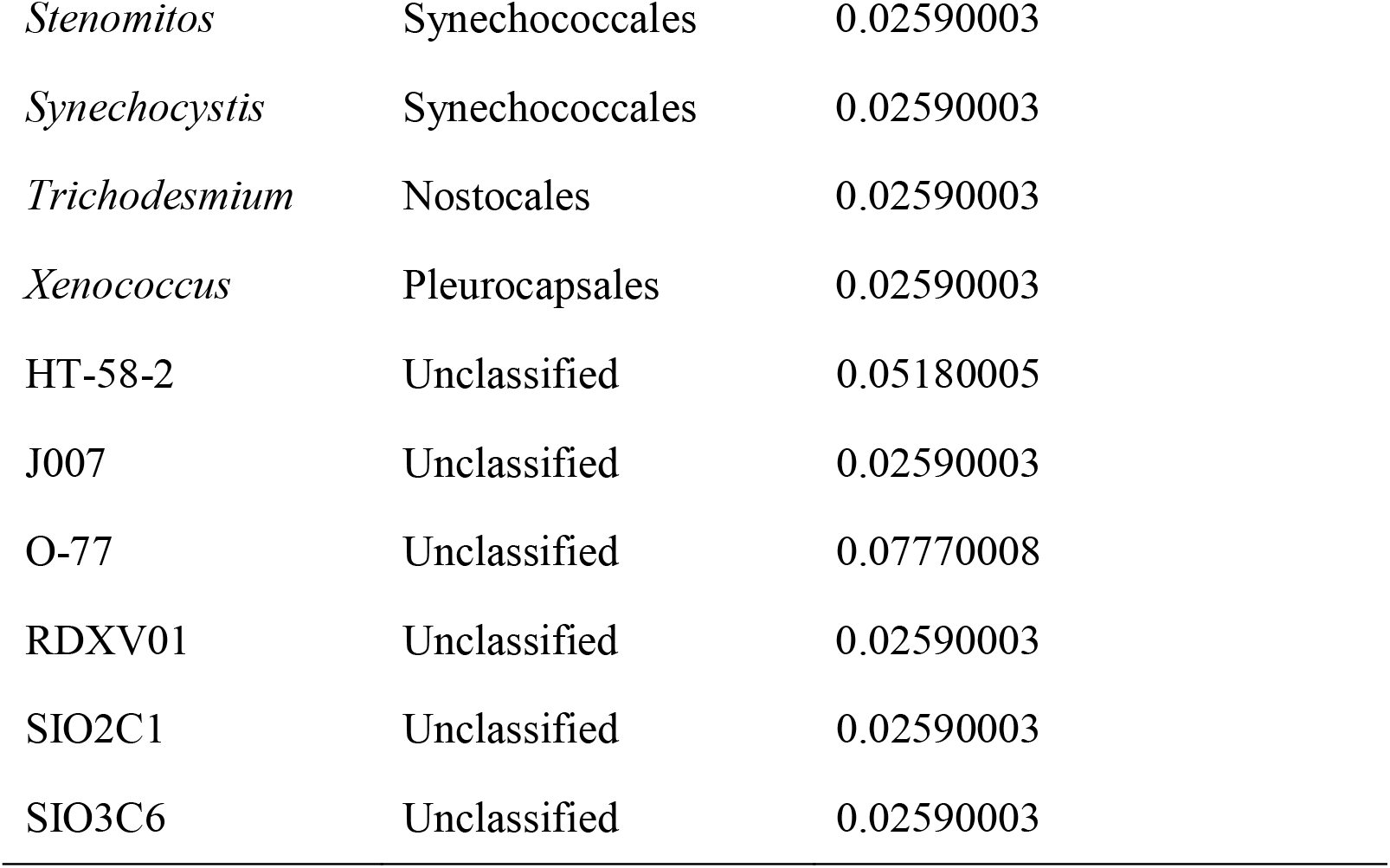
List of all cyanobacterial genera in the assembled genome of the stromatolites from San Felipe Creek in the Anza Borrego Desert (45 genera classified into 7 orders and 6 unclassified cyanobacteria).

Furthermore, *Rivularia* plays an important role in the calcification process of the stromatolite formation. *Rivularia* forms multilayered colonies, with pronounced seasonal growth and calcification patterns in freshwater habitats (Pentecost 1987). *Rivularia* was the main component of stromatolite-like formations observed in non-perennial calcareous streams in Spain (Aboal 1989) and was recorded in the stromatolite cyanobacterial community of San Felipe Creek with 0.07% abundance. In another segments of San Felipe Creek we collected large dry mats with filamentous cyanobacterium *Limnoraphis* (order Oscillatoriates) which formed wide calcium impregnated sheaths, closely resembling in size and structure the calcite tubes illustrated by Buchheim (1995) in stromatolites from a stream in the Anza Borrego Desert. Although *Limnoraphis* was not recorded in the metagenome of the studied stromatolites, morphologically similar genera were detected (e.g. *Lyngbya, Moorea, Okeania* (Table 2).

Furthermore, nostocacean cyanobacteria produce extensive extracellular mucilage and thick-walled akinetes, which protect the cells from desiccation (Komárek and Johansen 2015). Akinetes can be dormant for a long time and germinate immediately upon favorable conditions and water availability (Agrawal 2009). Major genera give insight into what key functions are being carried out for the stromatolite’s survival and maintenance.

We compared the metagenomes of freshwater stromatolites from Anza Borrego Desert with available data for hypersaline stromatolites (Babilonia et al. 2018). As expected, due to contrasting salinities and hydrological conditions, stromatolite microbial communities were taxonomically very different. The high abundance of Cyanobacteria differentiated the freshwater riverine stromatolite community in Anza Borrego Desert from marine stromatolites in South Shark Bay (SRX3553194), Central Shark Bay (SRX3553203), and North Shark Bay (SRX3553197). Proteobacteria and Actinobacteriota were the most abundant organisms in marine stromatolite sites, which opposed what was seen in Anza Borrego (Figure 4A). Since Cyanobacteria did not dominate their metagenomes, there was more evenness at the genus level South Shark Bay (Shannon Index = 8.07), Central Shark Bay (Shannon Index = 7.95), and North Shark Bay (Shannon Index 8.12)), with most falling below the 2% abundance threshold for all other sites (Figure 4C-D).

**Figure 4.**
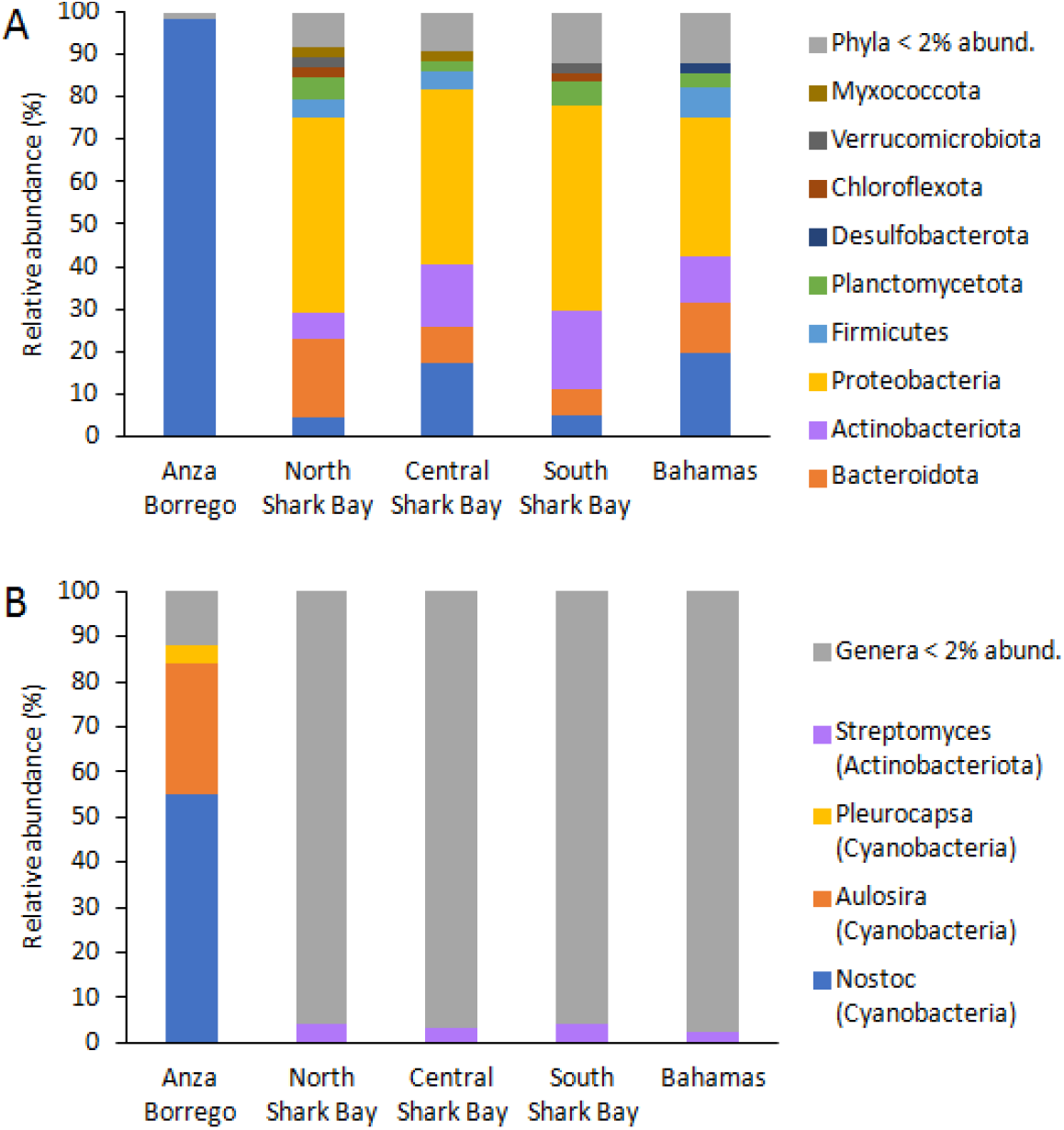
Taxonomic composition of the microbial community in the freshwater stromatolites of the San Felipe Creek in Anza Borrego compared to marine stromatolites in Shark Bay (Bibilonia et al., 2018) at phylum (A) and genus (B) level. Data for the complete prokaryote microbial community for the raw sequence reads are presented.

The taxonomic composition of Anza Borrego stromatolites was similar to freshwater stromatolites from pools in Cuatros Cienegas Basin (Breitbart et al. 2008) and Ruidera, Spain (Santos et al. 2010) which were also dominated by cyanobacteria (74% and 54% respectively). Unfortunately, we were not able to compare in detail the three freshwater stromatolite communities due to discrepancies in the molecular methods. The criterion for comparison was whole metagenomes above 2G bases, which was not satisfied by the currently available freshwater stromatolite sequence data.

### Gene Prediction and Functional Annotation

Insight into the ecological function of the microbial community was gleaned by annotating the genome using PGAP and functionally classifying the predicted gene sequences using KEGG. Gene prediction using PGAP resulted in community level annotation of ∼33,000 protein coding genes (Table 1), including 7 complete 5S rRNA, and 16 16S rRNA, and 34 23S partial rRNA genes. Of the protein coding sequences putatively identified, 11,732 or approximately 37% were functionally classified using GHOSTKOALA, with the largest functional category (10%) being affiliated with signaling and cellular processes. Additionally, a large number of genes were linked to genetic information processing (9%), carbohydrate metabolism (9%), and environmental information processing (9%) (Figure 5).

**Figure 5.**
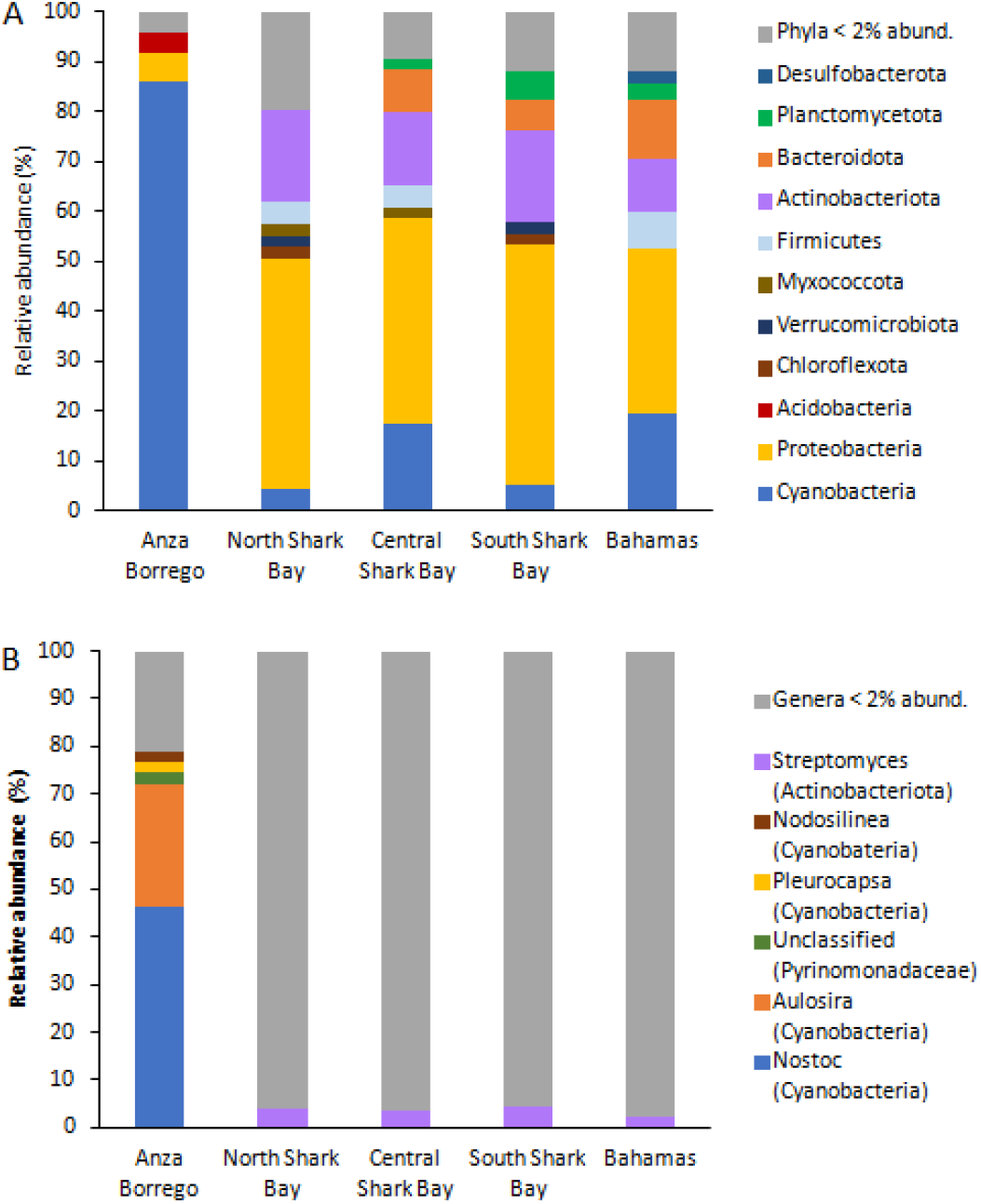
Taxonomic composition of the microbial community in the freshwater stromatolites of the San Felipe Creek in Anza Borrego compared to marine stromatolites in Shark Bay (Bibilonia et al., 2018) at phylum (A) and genus level (B). Data for the complete prokaryote microbial community for the assembled metagenome are presented.

**Figure 6.**
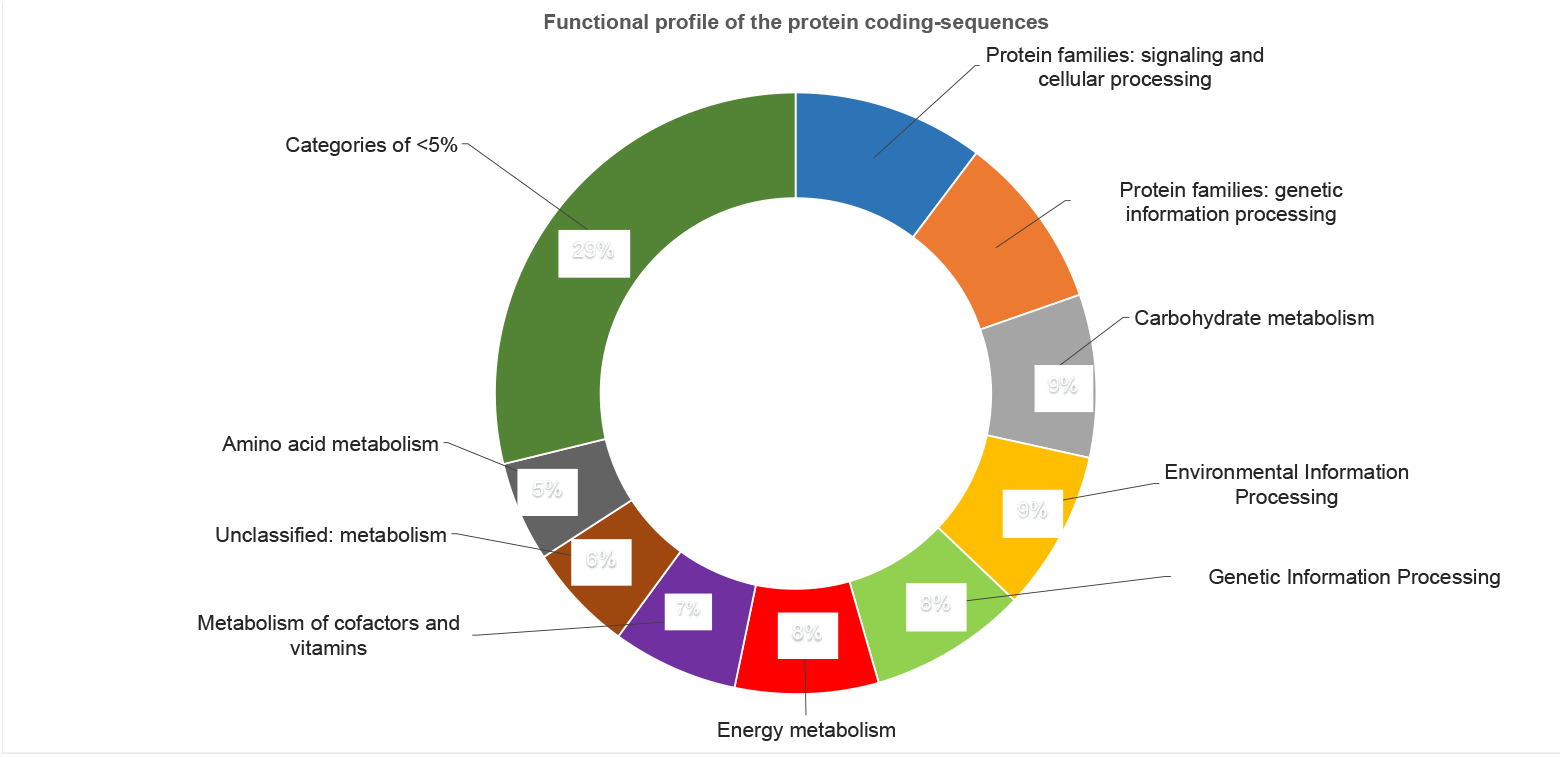
Functional profiles of protein-coding sequences. Approximately 40% of the annotated proteins were identified and classified using GhostKoala against prokaryotic, eukaryotic and viral databases. Across all functional categories, signaling and cellular processing protein families (10%) were the most represented followed by those involved genetic information processing (9%), carbohydrate metabolism (9%) and environmental information processing (9%).

### Viruses

Of significance 173 genes showed similarity to viral coding sequences indicating a possible functional role of viruses in the formation of stromatolites as has been previously documented (Desnues et al., 2008). The KEGG similarity scores of the viral sequences range from 27-605. Altogether 112 different viruses were detected. Among those detected, sequences showing homology to genes from the double stranded DNA Mycobacterium phage (∼20%) were the most prevalent followed by genes from *Nodularia* phage (∼10%). The viruses present in the stromatolites, particularly *Nodularia* phage vB_NpeS-2AV2 are likely to impact community population dynamics and activity. For example, viral lysis of nitrogen-fixing cyanobacterium such as *Nodularia* releases nutrients, importantly dissolved organic and inorganic nitrogen which can be scavenged by other microbes for the synthesis of proteins and nucleic acids. Nitrogen can also affect the pH of the microenvironment to ultimately influence microbial growth.

### Biofilm and Quorum Sensing Proteins

The stromatolites in the Anza Borrego desert reside in a brackish environment where the resident microbes must be able to adapt to the harsh conditions of extreme heat, high UV light, desiccation and fluctuating solubilization/precipitation and hydration/evaporation. To persist and survive in these conditions the highly adapted and evolved microbes employ active defense strategies, some of which involve the formation of biofilms and quorum sensing (QS). There are 154 genes with 35 unique functions (Supplementary Table S3) related to bacterial biofilms (KO 05111, 02025, and 02026) present in the stromatolite metagenome. These genes suggest at times the microbial community may be encased in an extracellular polymeric matrix composed primarily of polysaccharides (*PYG, bcsA, glgA, glgC, pslH, wecB*), and possibly lipids, proteins (*trpE, trpG, cysE*), and nucleic acids. Apart from desiccation protection, this biofilm is also likely to contribute to the formation of the stromatolites, promoting lithification by facilitating sediment trapping, binding, and/or precipitation as a function of alterations in water chemistry brought about by microbial growth and metabolic activities (Noffke and Awramik, 2013). As noted previously (Vanada et al., 2022) and evidenced here, the formation of biofilms is a complex and dynamic process, controlled by regulatory networks that include transcriptional (*crp, hfq*) and post transcriptional regulators such as two component chemosensory (*wspA, wspB, wspF*) and twitching motility systems (*pilH, pilH, pilG*), alternative sigma factors (rpoS), second messengers (*cpdA, ADCY*), export or secretory systems (*exoP, vspO, impJ, vase, gspE, wza, gfcE*), and surface proteins (*BapA*), among others. The spatial complexity of the biofilms formed by the stromatolite community also appears to be tightly controlled by quorum sensing (QS) molecules including Ni^2+^ and Cd^2+^, anthranilate, spermidine, putrescine, and rhamnolipids. Genes implicating some of these molecules include permeases for the transport for Ni^2+^ and Cd^2+^ (*nikA*) that decrease the expression of genes responsible for acyl homoserine lactone (AHL) QS at the transcriptional level to inhibit biofilm formation (Vega et al., 2014); genes involved in the synthesis anthranilate (*trypE, trpG*), a transcriptional regulator that also attenuates biofilm formation; a gene encoding the binding protein for the transport of spermidine and putrescine (ABC.SP.S) which facilitate the efflux of AHLs to increase biofilm deposition; and genes involved in the production of rhamnolipids (*rhlB, rfbF, and rhlC*) that inhibit the synthesis of, and disrupt preformed biofilms (Sidrim et al., 2020; Malakar er al., 2021).

Although many bacteria can survive as single cells, microbes responsible for the formation of stromatolites reside in a biofilm cooperative where the chemical communication and quorum sensing systems they employ enables them to synchronize their physiological and transcriptional processes. As a cooperative they can strategically manipulate the environment to enhance survival possibilities (Goo et al., 2015; Khider et al., 2019). Several quorum sensing systems (QS) used to manipulate microbial activities based on cell densities were identified in the San Felipe creek stromatolites. More than 400 genes encoding 34 different proteins (Supplementary Table S4, KO 02024) were annotated related to the production, detection, and accumulation of QS signaling molecules and the physiological response patterns. While neither acyl homoserine lactone synthase (AHLs) genes or those encoding synthases that produce other autoinducers were detected, sequences showing homology to *LuxR* transcriptional regulators that employ acyl-homoserine lactones (AHL) as signal molecules, the QS quenching N-acyl homoserine lactone hydrolase (*aiiA*), and a long-chain acyl-CoA synthetase (acsl) were identified. Hence, it is likely that acyl homoserine lactones (AHL) facilitate the cell-to-cell communication between microbes in the stromatolites necessary to orchestrate communal activities such as biofilm formation, competence, aggregation, and virulence factor production.

### Nitrogen cycling

Nitrogen cycling is likely to play a fundamental role in mediating both the structure of the stromatolite microbial community and mineral precipitation/dissolution. Metagenome annotations revealed complete pathways for nitrogen fixation and assimilatory nitrate reduction, and partial pathways for dissimilatory nitrate reduction, denitrification, and nitrification (Supplementary Table S5). The N demand of organisms residing in the stromatolites is likely to be met by cyanobacteria. Detection of seven unique molybdenum- and nine iron containing nitrogenases (*nifD, nifH, nifK*) in the metagenome indicates a variety of N2-fixing cyanobacterial genera (*Nodularia, Nostoc, Calothrix, Trichormus, Anabaena, Tolypothrix)* and, *Chroococcidiopsis* contribute to the bioavailability of N in the oligotrophic environment of the stromatolites. A multitude of other genes (*nifX, nifB, nifZ, nifT, nifEN, nifN, nifU, nifE, nifV, nifS*) that contribute to the efficient conversion of atmospheric nitrogen to ammonia were also detected and implicate additional filamentous (*Cylindrospermum, Rivularia, Fischerella*) and coccoid (*Chrondrocystis, Stanieria*) nitrogen fixing cyanobacteria. Sequence data also indicates Azotobacter, one of a limited number of bacterium known to fix nitrogen, is also present in the stromatolites. Annotation revealed ureolysis which generates ammonia and carbonic acid, is another viable source of N for the microbial community. Urease (*UreA, UreB, UreC*) and accessory protein (*UreD, UreE, UreF, UreG*) homologs were identified in several of the same N2-fixing cyanobacterial lineages (Supplementary Table S5). Although not detected here, possibly due to lack of sequencing depth, urea is a nitrogen source for a variety of fungi and bacteria in addition to cyanobacteria. Apart from serving as a N-source, the hydrolysis of urea in the stromatolites is particularly noteworthy. When urea is hydrolyzed, it produces ammonia which is further hydrolyzed generating OH^-^ ions and increases the pH of the microenvironment. Additionally, carbonic acid is produced and dissociates yielding bicarbonate which increases the dissolved inorganic carbon (DIC). These reactions combined increase the overall alkalinity and in the presence of calcium, favors calcium carbonate precipitation which likely contributes to the formation of stromatolites.

Denitrification provides energy and supports microbial growth when oxygen is limiting, and nitrate and organic carbon are plentiful. Functional markers for denitrification were identified in the metagenome including nine unique nitrate reductase (*narB, narG, narZ, nxrA*) and 10 different nitrite reductase genes (*nirA*). These homologs implicate cyanobacteria as the primary denitrifiers in the microbial community (Supplementary Table S5). Denitrification imparts additional flexibility in terms of microbial growth strategies but also impacts biomineralization by consuming H^+^ and producing CO_2_ which further generates carbonate ions. Nitrogen can also be obtained by the ammonification or the degradation of amino acids which typically occurs under aerobic conditions and involves various lyases (aspartase, histidase, etc.) and hydrolases (asparaginase, glutaminase, amidase, and arginase). Evidence from the annotated stromatolite metagenome suggests several cyanobacteria utilize amino acids for metabolic purposes. When amino acids are used as a nitrogen source, the excess ammonium, because of its toxicity, is probably secreted. When secreted, it becomes available for microbially-mediated nitrification using ammonia monooxygenase (AMO) and hydroxylamine oxidoreductase (HAO). Genes encoding these enzymes, however, were not detected, and the ammonia in the microenvironment may be hydrolyzed and once again, promote the precipitation of carbonate biominerals. Taken together, the genomic evidence supports the contention that the diversity and abundance of cyanobacteria in San Filipe stromatolites play a fundamental role in modulating total nitrogen which strongly influences microbial growth, but also impacts water chemistry and mineral precipitation.

### Metals and Other Key Nutrients

Stromatolite ecosystems provide a diverse array of microniches where different functional guilds can develop complex metabolite exchange with the substrate that ensure their survival under extreme conditions (Paerl and Yannarell, 2010). Studied stromatolite metagenomes reveal several genes that might enable a specialized and unique group of endolithic cyanobacteria including *Chroococcidiopsis, Hyella, Myxosarcina*, and *Pleurocapsa* (Komárek and Johansen, 2015) to derive metals and important nutrients from rocks. Some of these genes are involved in the process of chelation and transport of metals while other genes encode specialized enzymes that can break down minerals in rock and release nutrients such as phosphate, sulfate, and nitrogen (Gadd, 2010). Cyanobacteria boring into limestone substrata may cause biogenic destruction of lithified stromatolite. Similar endolithic, photosynthetic cyanobacteria were identified in hot spring-formed ancient and recent travertine deposits in Yellowstone National Park, USA (Norris and Castenholz 2006).

Furthermore, cyanobacteria in the stromatolites from San Felipe Creek possess copper oxidases (*MCO*) and Zip (*ZIP*), Nramp (*mntH*) and other metal transporters (Supplementary Table S6) that are likely responsible for the uptake of metal ions from the environment. With iron often limiting in freshwater streams, siderophore receptor (*FEV*.*OM*) and transport genes (*ABC*.*FEV*.*S*) present in the metagenome suggest these organic compounds are used to chelate iron and other metal ions, possibly from the rock and the desert stream, when water is available (Supplementary Table S6). Phosphorus is another important nutrient that is often limiting in freshwater streams, and hence the detecting numerous phosphatase genes (*PhoD*, in particular) is not surprising and suggests the cyanobacteria are able to release phosphorus from organic material in the rocks and sediments. Cyanobacteria in the stromatolites are likely to secrete H^+^, organic acids, and other metabolites which can solubilize rock minerals and nutrients that can then be absorbed for their growth and metabolism.

### Desiccation strategies

Cyanobacteria can efficiently exploit limiting and/or pulsating resources and adapt to fluctuating environmental conditions including variations in temperature, light, salinity, and moisture.

Activity and survival in stress laden habitats such as that of the Anza Borrego desert where during the study period (January 2019 to January 2023) monthly highest max temperature ranged from 72° F to 121° F, and total annual precipitation ranged from 2.76 in for 2022 to 9.25 in for 2019, is dependent upon key biomolecules that provide protection against adverse conditions. Cyanobacteria tolerate high and low extremes of temperature by the ability to synthesize protective pigments is particularly important during the spring and summer months. Metagenomic signatures were found for scytonemin (*ScyC, ScyA, ScyF, EboC*), carotenoids (*crtO, Ocp, Dxr, Dxs, Pds,Psy,CrtH, Zds, Ggr*), and the synthesis of potential microsporine-like amino acids (DHQ synthase, sedohepulose 7-phosphate cyclase) (Supplementary Table S7).

As a sunscreen, scytonemin accumulates in cyanobacteria residing in highly exposed habitats to guard against photoinhibition and UV radiation damage, while carotenoids are amassed and offer photoprotection and quenching of free radicals resulting from UV exposure (Wada et al., 2013). Microsporine-like amino acids may be accrued and function as UV-A light absorbing agents (Wada et al., 2013). Signature genes essential to the biosynthesis of phycocyanin present in the metagenome span ten species and include *cpcA, cpcB, apcA, cpcT/CpeT* (Supplementary Table S7). Phycocyanin is an accessory pigment that enables cyanobacteria to survive under low light conditions as might be expected when communities reside in stromatolite zones beneath layers of calcium carbonate.

Members of the stromatolite community possess different morphological and physiological strategies to cope with desiccation and hydric stress that reflect the San Felipe creek as an intermittent desert stream. Hydric pressures imposed by the shortage of water in the creek bed is reflected more readily in transcriptomic data, and likely to lead to reduced growth rates, decreased reproductive success, and greater susceptibility to viral infection, toxins, and other stressors. The metagenomic data, however, revealed various genes involved in microalgal desiccation tolerance, including those encoding aquaporins (*Aqp*), chaperones (*DnaK, DnaJ, GroEL*), antioxidant enzymes (*SOD, GPx, CAT*), enzymes responsible for the synthesis of trehalose (*TreY, TreZ*) sucrose (*Sus*), and proteins required for the synthesis of polyamines such as spermidine (*SpdS*), (Supplementary Table 8). These genes play a critical role in protecting cyanobacteria against desiccation-induced damage by maintaining cellular hydration, stabilizing proteins, scavenging reactive oxygen species, maintaining osmotic balance, and regulating gene expression. Desiccation tolerance is strongly associated with the synthesis of disaccharides such as trehalose and sucrose and extracellular polysaccharide (EPS). In addition to being an osmoprotectant, trehalose stabilizes proteins and cell membranes by substituting for water and creating a hydrophilic shield around biomolecules, while sucrose serves as a compatible solute.

Some cyanobacteria undergo morphological changes in response to dehydration. The formation of akinetes by members of the stromatolite community is implicated by the presence of *HetR* and *HetP* genes in the metagenome. *HetR* is a transcriptional regulator that controls the expression of genes involved in the differentiation of specialized dormant cells that enable some cyanobacterial to survive under adverse environmental conditions. *HetP* is thought to regulate hetR and other genes that participate in akinete cell wall synthesis (Leganes et al., 1994). Other akinete specific genes including *akinA, akinB*, and *hrmA*, however, were not detected, perhaps due to inadequate depth of sequencing. Select cyanobacteria in the stromatolites of the San Felipe Creek are also likely to enter a form of dormancy known as anhydrobiosis where they can lose up to 95% of their water content and remain in a suspended state of animation during which time metabolic process are slowed down or stopped for the purpose of self-preservation under extreme conditions or, in this case, for extended dry periods, until water becomes available again. Anhydrobiotes species present in the stromatolites include *Nostoc commune* (Rajeev et al., 2013), *Nostoc punctiforme* (Argueta et al., 2006), *Chroococcidiopsis sp*. (Billi et al., 2000), and *Anabaena sp*.(Rajeev et al., 2013). Although there are no universal biomarker genes, these cyanobacteria and other anhydrobiotes undergo morphological changes and survive long dry periods and hydric stress without sustaining irreparable damage to cellular structures using a variety of molecular and physiological mechanisms. In the case of *Nostoc commune*, cells become shorter in length and thickness, with cell walls that are more electron dense (Olsson-Francis et al., 2017). *Chroococcidiopsis sp*. cells become smaller, more spherical, and exhibit tightly packed thylakoid membranes (Billi et al., 2000). In both instances, cells return to the original size and share upon rehydration.

## Conclusions

Metagenomic studies of living stromatolites is a powerful tool for addressing fundamental questions that span diverse fields from microbiology, geochemistry, evolution, phycology, astrobiology, to paleoecology. This study investigated the microbial community structure and functional characteristics of freshwater stromatolites in the harsh open canopy environment of the ephemeral San Felipe Creek in the Anza Borrego desert. Molecular signals revealed the adaptive strategies employed by the microbial community to survive ecosystem where they are constantly exposed to high UV light, desiccation, and fluctuating hypersaline waters whose cellular and metabolic process impact mineral products. An emerging trend is that the microbial and metabolic processes have an impact on mineral products, as evidenced by the community composition which is predominately made up of cyanobacteria. The stromatolites in this environment are formed through the rapid and plentiful growth of blue green algae that metabolically contribute to the precipitation of minerals and whose biofilms trap fine sediments. This results in periodic layering of calcite, detrital stream particles, and an organic component consisting largely of cyanobacteria.

## Data availability

The assembled metagenome has been submitted to NCBI and is accessible via BioProject: PRJNA967693.This Whole Genome Shotgun project has been deposited at DDBJ/ENA/GenBank under the accession JASEJY000000000. The version described in this paper is version JASEJY010000000. Scripts and code for all analyses performed can be accessed via the project’s GitHub page, accessible at https://github.com/j-archambeau/Stromatolite_Metagenomics

## Acknowledgments

AS and BR thank the mentors of the Summer 2022 CSUSM NSF REU scholars for their continued support and encouragement for student-led research. We thank Douglas B. Read Jr. for stromatolite images presented in Figure 1.

## Funding

This work was funded by NSF-REU: 1852189 to PI Betsy Read, co-PI Sethuraman, and senior personnel Rosalina Stancheva and Elinne Becket, and was conducted over Summer 2022 by CSUSM NSF REU students whose experiences were chronicled at www.csusmbioreu.weebly.com. This research includes calculations carried out on HPC resources supported in part by the National Science Foundation through major research instrumentation grant number 1625061 and by the US Army Research Laboratory under contract number W911NF-16-2-0189.

## Notes

### Competing Interest Statement

The authors have declared no competing interest.

