## Supplemental Tables for "Characterizing the microbial metagenome of calcareous stromatolite formations in the San Felipe Creek in Anza Borrego Desert"

Table S1: Metagenome Assemblies Statistics. DNA was isolated from four stromatolites samples and libraries were prepared, sequenced, and assembled.

| Parameter/Statistic | Metagenome 1 | Metagenome 2 | Metagenome 3 | Metagenome 4 |
| --- | --- | --- | --- | --- |
| Genome Assembly |  |  |  |  |
| #Contig (≥ 0 bp) | 886 | 2763 | 1801 | 460 |
| #Contig (≥1000 bp) | 96 | 214 | 125 | 45 |
| Total Length | 684,520 | 1,947,576 | 1,283,752 | 363,808 |
| Total Length (≥1000 bp) | 198,852 | 352,372 | 243,579 | 108,066 |
| Largest Contig | 65,283 | 49,277 | 65,402 | 49,277 |
| % G+C | 45.09 | 42.61 | 42.33 | 44.86 |
| N50 | 670 | 649 | 645 | 675 |
| L50 | 282 | 1025 | 651 | 138 |
| Genome Annotation |  |  |  |  |
| Genes (total) | 11,216 | 20,880 | 16,709 | 8,073 |
| CDSs (with protein homology) | 10,584 | 19,994 | 15,974 | 7,384 |
| Ab-initio predicted | 417 | 542 | 427 | 517 |
| Genes (RNA) | 128 | 164 | 156 | 113 |
| rRNAs | 6, 21, 55 (5S, 16S, 23S) | 5, 28, 62 (5S, 16S, 23S) | 6, 27, 58 (5S, 16S, 23S) | 4, 27, 47 (5S, 16S, 23S) |
| tRNAs | 43 | 66 | 61 | 35 |
| ncRNAs | 3 | 3 | 4 | 0 |
| Pseudo Genes (total) | 87 | 180 | 152 | 59 |

Table S2: Bins based on the Assembled Composite Metagenome

**Bin 1**

Predicted organism: Anabaena cylindrica (taxid = 1165, rank = species, lineage = Bacteria; Cyanobacteria; Nostocales; Nostocaceae; Anabaena)
Submitted organism has type: No
Status: INCONCLUSIVE
Confidence: LOW
90.899 ( 7.5	.0)	504428 assembly	2835 Anabaena cylindrica PCC 7122 (GCA_000317695.1, ASM31769v1)

**Bin 2**

Predicted organism: (none)
Submitted organism has type: No
Status: INCONCLUSIVE
Confidence: LOW

**Bin 3**

Predicted organism: (none)
Submitted organism has type: No
Status: INCONCLUSIVE
Confidence: LOW

**Bin 4**

Predicted organism: Halomonas lysinitropha (taxid = 2607506, rank = species, lineage = Bacteria; Proteobacteria; Gammaproteobacteria; Oceanospirillales; Halomonadaceae; Halomonas)
Submitted organism has type: No
Status: INCONCLUSIVE
Confidence: LOW
91.387 (31.8	.1) 6943128 assembly	2620 Halomonas montanilacus (GCA_003336675.1, ASM333667v1)

**Bin 5**

Predicted organism: (none)
Submitted organism has type: No
Status: INCONCLUSIVE
Confidence: LOW

**Bin 6**

Predicted organism: (none)
Submitted organism has type: No
Status: INCONCLUSIVE
Confidence: LOW

**Bin 7**

Predicted organism: (none)
Submitted organism has type: No
Status: INCONCLUSIVE
Confidence: LOW

**Bin 8**

Predicted organism: (none)
Submitted organism has type: No
Status: INCONCLUSIVE
Confidence: LOW
86.508 (19.8 37.1) 20828 assembly 3344689 Nostoc punctiforme PCC 73102 (GCA_000020025.1, ASM2002v1)

**Bin 9**

Predicted organism: Tolypothrix tenuis (taxid = 457083, rank = species, lineage = Bacteria; Cyanobacteria; Nostocales; Tolypothrichaceae; Tolypothrix)
Submitted organism has type: No
Status: INCONCLUSIVE
Confidence: LOW
92.299 (19.0	.0) 5101288 assembly	935 Tolypothrix tenuis PCC 7101 (GCA_002368295.1, ASM236829v1)

**Bin 10**

Predicted organism: Nostoc punctiforme (taxid = 272131, rank = species, lineage = Bacteria; Cyanobacteria; Nostocales; Nostocaceae; Nostoc)
Submitted organism has type: No
Status: INCONCLUSIVE
Confidence: LOW
88.525 (45.0	.0)	20828 assembly	4366 Nostoc punctiforme PCC 73102 (GCA_000020025.1, ASM2002v1)

**Bin 11**

Predicted organism: (none)
Submitted organism has type: No
Status: INCONCLUSIVE
Confidence: LOW

**Bin 12**

Submitted organism has type: No
Status: INCONCLUSIVE
Confidence: LOW
82.808 (62.8	.0)	792958 assembly	762 Rubidibacter lacunae KORDI 51-2 (GCA_000473895.1, KS51_v1

**Bin 13**

Predicted organism: (none)
Submitted organism has type: No
Status: INCONCLUSIVE
Confidence: LOW

**Bin 14**

Predicted organism: (none)
Submitted organism has type: No
Status: INCONCLUSIVE
Confidence: LOW

**Bin 15**

Predicted organism: (none)
Submitted organism has type: No
Status: INCONCLUSIVE
Confidence: LOW

**Bin 16**

Predicted organism: (none)
Submitted organism has type: No
Status: INCONCLUSIVE
Confidence: LOW

**Bin 17**

Predicted organism: (none)
Submitted organism has type: No
Status: INCONCLUSIVE
Confidence: LOW

**Bin 18**

Predicted organism: (none)
Submitted organism has type: No
Status: INCONCLUSIVE
Confidence: LOW

**Bin 19**

Predicted organism: Sphingobium indicum (taxid = 332055, rank = species, lineage = Bacteria; Proteobacteria; Alphaproteobacteria; Sphingomonadales; Sphingomonadaceae; Sphingobium)
Submitted organism has type: No
Status: INCONCLUSIVE
Confidence: LOW
93.976 ( .4	.0) 6687368 assembly	0 Rhodobacter capsulatus (GCA_003254295.1, ASM325429v1)

**Bin 20**

Predicted organism: (none)
Submitted organism has type: No
Status: INCONCLUSIVE
Confidence: LOW

**Bin 21**

Predicted organism: (none)
Submitted organism has type: No
Status: INCONCLUSIVE
Confidence: LOW

**Bin 22**

Predicted organism: (none)
Submitted organism has type: No
Status: INCONCLUSIVE
Confidence: LOW

**Bin 23**

Predicted organism: Halomonas anticariensis (taxid = 258591, rank = species, lineage = Bacteria; Proteobacteria; Gammaproteobacteria; Oceanospirillales; Halomonadaceae; Halomonas)
Submitted organism has type: No
Status: INCONCLUSIVE
Confidence: LOW
88.252 ( 1.4	.0)	778038 assembly	0 Halomonas anticariensis FP35 = DSM 16096 (GCA_000428505.1, ASM42850v1)

**Bin 24**

Predicted organism: (none)
Submitted organism has type: No
Status: INCONCLUSIVE
Confidence: LOW
96.875 ( .0	.0) 8176428 assembly	0 Chlorogloeopsis fritschii PCC 6912 (GCA_003990575.1, ASM399057v1)

**Bin 25**

Predicted organism: (none)
Submitted organism has type: No
Status: INCONCLUSIVE
Confidence: LOW

**Bin 26**

Predicted organism: (none)
Submitted organism has type: No
Status: INCONCLUSIVE
Confidence: LOW

**Bin 27**

Predicted organism: Nostoc punctiforme (taxid = 272131, rank = species, lineage = Bacteria; Cyanobacteria; Nostocales; Nostocaceae; Nostoc)
Submitted organism has type: No
Status: INCONCLUSIVE
Confidence: LOW
90.452 (44.9	.0)	20828 assembly	2168 Nostoc punctiforme PCC 73102 (GCA_000020025.1, ASM2002v1)

**Bin 28**

Predicted organism: (none)
Submitted organism has type: No
Status: INCONCLUSIVE
Confidence: LOW
90.278 (45.6	.0) 5101288 assembly	3877 Tolypothrix tenuis PCC 7101 (GCA_002368295.1, ASM236829v1)

**Bin 29**

Predicted organism: (none)
Submitted organism has type: No
Status: INCONCLUSIVE
Confidence: LOW

**Bin 30**

Predicted organism: (none)
Submitted organism has type: No
Status: INCONCLUSIVE
Confidence: LOW

**Bin 31**

Predicted organism: (none)
Submitted organism has type: No
Status: INCONCLUSIVE
Confidence: LOW

**Bin 32**

Predicted organism: (none)
Submitted organism has type: No
Status: INCONCLUSIVE
Confidence: LOW

**Bin 33**

Predicted organism: Nostoc punctiforme (taxid = 272131, rank = species, lineage = Bacteria; Cyanobacteria; Nostocales; Nostocaceae; Nostoc)
Submitted organism has type: No
Status: INCONCLUSIVE
Confidence: LOW
86.646 (16.8	.0)	20828 assembly	1273 Nostoc punctiforme PCC 73102 (GCA_000020025.1, ASM2002v1)

**Bin 34**

Predicted organism: (none)
Submitted organism has type: No
Status: INCONCLUSIVE
Confidence: LOW

**Bin 35**

Predicted organism: (none)
Submitted organism has type: No
Status: INCONCLUSIVE
Confidence: LOW

**Bin 36**

Predicted organism: Nostoc punctiforme (taxid = 272131, rank = species, lineage = Bacteria; Cyanobacteria; Nostocales; Nostocaceae; Nostoc)
Submitted organism has type: No
Status: INCONCLUSIVE
Confidence: LOW
88.489 (77.3	.0)	20828 assembly	738 Nostoc punctiforme PCC 73102 (GCA_000020025.1, ASM2002v1)

**Bin 37**

Predicted organism: (none)
Submitted organism has type: No
Status: INCONCLUSIVE
Confidence: LOW

**Bin 38**

Predicted organism: Tolypothrix tenuis (taxid = 457083, rank = species, lineage = Bacteria; Cyanobacteria; Nostocales; Tolypothrichaceae; Tolypothrix)
Submitted organism has type: No
Status: INCONCLUSIVE
Confidence: LOW
83.556 (47.6	.0) 5101288 assembly	1198 Tolypothrix tenuis PCC 7101 (GCA_002368295.1, ASM236829v1)

**Bin 39**

Predicted organism: (none)
Submitted organism has type: No
Status: INCONCLUSIVE
Confidence: LOW

**Bin 40**

Predicted organism: (none)
Submitted organism has type: No
Status: INCONCLUSIVE
Confidence: LOW

**Bin 41**

Predicted organism: Nostoc punctiforme (taxid = 272131, rank = species, lineage = Bacteria; Cyanobacteria; Nostocales; Nostocaceae; Nostoc)
Submitted organism has type: No
Status: INCONCLUSIVE
Confidence: LOW
91.195 (17.9	.0)	20828 assembly	579 Nostoc punctiforme PCC 73102 (GCA_000020025.1, ASM2002v1)

**Bin 42**

Predicted organism: Paracraurococcus ruber (taxid = 77675, rank = species, lineage = Bacteria; Proteobacteria; Alphaproteobacteria; Rhodospirillales; Acetobacteraceae; Paracraurococcus)
Submitted organism has type: No
Status: INCONCLUSIVE
Confidence: LOW
76.695 ( 1.3	.0) 8757698 assembly	944 Paracraurococcus ruber (GCA_004353985.1, ASM435398v1)

**Bin 43**

Predicted organism: (none)
Submitted organism has type: No
Status: INCONCLUSIVE
Confidence: LOW

**Bin 44**

Predicted organism: (none)
Submitted organism has type: No
Status: INCONCLUSIVE
Confidence: LOW

**Bin 45**

Predicted organism: (none)
Submitted organism has type: No
Status: INCONCLUSIVE
Confidence: LOW

**Bin 46**

Predicted organism: (none)
Submitted organism has type: No
Status: INCONCLUSIVE
Confidence: LOW
83.913 (80.2	.0) 5101288 assembly	0 Tolypothrix tenuis PCC 7101 (GCA_002368295.1, ASM236829v1)

**Bin 47**

Predicted organism: Nostoc punctiforme (taxid = 272131, rank = species, lineage = Bacteria; Cyanobacteria; Nostocales; Nostocaceae; Nostoc)
Submitted organism has type: No
Status: INCONCLUSIVE
Confidence: LOW
74.349 (46.8	.0)	20828 assembly	1727 Nostoc punctiforme PCC 73102 (GCA_000020025.1, ASM2002v1)

Bin 48

Predicted organism: Nostoc punctiforme (taxid = 272131, rank = species, lineage = Bacteria; Cyanobacteria; Nostocales; Nostocaceae; Nostoc)
Submitted organism has type: No
Status: INCONCLUSIVE
Confidence: LOW
93.177 (71.5	.0)	20828 assembly	3195 Nostoc punctiforme PCC 73102 (GCA_000020025.1, ASM2002v1)

**Bin 49**

Predicted organism: (none)
Submitted organism has type: No
Status: INCONCLUSIVE
Confidence: LOW

**Bin 50**

Predicted organism: (none)
Submitted organism has type: No
Status: INCONCLUSIVE
Confidence: LOW
87.526 (54.4	.1)	20828 assembly	4742 Nostoc punctiforme PCC 73102 (GCA_000020025.1, ASM2002v1)

**Bin 51**

Predicted organism: (none)
Submitted organism has type: No
Status: INCONCLUSIVE
Confidence: LOW

**Bin 52**

Predicted organism: Anabaena cylindrica (taxid = 1165, rank = species, lineage = Bacteria; Cyanobacteria; Nostocales; Nostocaceae; Anabaena)
Submitted organism has type: No
Status: INCONCLUSIVE
Confidence: LOW
81.418 (27.9	.0) 5100948 assembly	0 Anabaena cylindrica PCC 7122 (GCA_002367955.1, ASM236795v1)

**Bin 53**

Predicted organism: (none)
Submitted organism has type: No
Status: INCONCLUSIVE
Confidence: LOW
97.403 ( .2	.0) 4984298 assembly	0 Rubrivirga marina (GCA_002283365.1, ASM228336v1)

**Bin 54**

Predicted organism: (none)
Submitted organism has type: No
Status: INCONCLUSIVE
Confidence: LOW

**Bin 55**

Predicted organism: (none)
Submitted organism has type: No
Status: INCONCLUSIVE
Confidence: LOW

**Bin 56**

Predicted organism: (none)
Submitted organism has type: No
Status: INCONCLUSIVE
Confidence: LOW

**Bin 57**

Predicted organism: (none)
Submitted organism has type: No
Status: INCONCLUSIVE
Confidence: LOW

**Bin 58**

Predicted organism: (none)
Submitted organism has type: No
Status: INCONCLUSIVE
Confidence: LOW

**Bin 59**

Predicted organism: (none)
Submitted organism has type: No
Status: INCONCLUSIVE
Confidence: LOW

**Bin 60**

Predicted organism: (none)
Submitted organism has type: No
Status: INCONCLUSIVE
Confidence: LOW

**Bin 61**

Predicted organism: (none)
Submitted organism has type: No
Status: INCONCLUSIVE
Confidence: LOW

**Bin 62**

Predicted organism: Nostoc punctiforme (taxid = 272131, rank = species, lineage = Bacteria; Cyanobacteria; Nostocales; Nostocaceae; Nostoc)
Submitted organism has type: No
Status: INCONCLUSIVE
Confidence: LOW
87.476 (86.4	.0)	20828 assembly	1062 Nostoc punctiforme PCC 73102 (GCA_000020025.1, ASM2002v1)

**Bin 63**

Predicted organism: Nostoc punctiforme (taxid = 272131, rank = species, lineage = Bacteria; Cyanobacteria; Nostocales; Nostocaceae; Nostoc)
Submitted organism has type: No
Status: INCONCLUSIVE
Confidence: LOW
89.366 ( 6.1	.0)	20828 assembly	3282 Nostoc punctiforme PCC 73102 (GCA_000020025.1, ASM2002v1)

**Bin 64**

Predicted organism: (none)
Submitted organism has type: No
Status: INCONCLUSIVE
Confidence: LOW

**Bin 65**

Predicted organism: (none)
Submitted organism has type: No
Status: INCONCLUSIVE
Confidence: LOW

**Bin 66**

Predicted organism: (none)
Submitted organism has type: No
Status: INCONCLUSIVE
Confidence: LOW

**Bin 67**

Predicted organism: (none)
Submitted organism has type: No
Status: INCONCLUSIVE
Confidence: LOW

**Bin 68**

Predicted organism: Tolypothrix tenuis (taxid = 457083, rank = species, lineage = Bacteria; Cyanobacteria; Nostocales; Tolypothrichaceae; Tolypothrix)
Submitted organism has type: No
Status: INCONCLUSIVE
Confidence: LOW
84.405 (95.4	.0) 5101288 assembly	1026 Tolypothrix tenuis PCC 7101 (GCA_002368295.1, ASM236829v1)

**Bin 69**

Predicted organism: (none)
Submitted organism has type: No
Status: INCONCLUSIVE
Confidence: LOW
87.185 (24.8 11.7) 20828 assembly 1029408 Nostoc punctiforme PCC 73102 (GCA_000020025.1, ASM2002v1)

**Bin 70**

Predicted organism: (none)
Submitted organism has type: No
Status: INCONCLUSIVE
Confidence: LOW

**Bin 71**

Predicted organism: (none)
Submitted organism has type: No
Status: INCONCLUSIVE
Confidence: LOW

**Bin 72**

Predicted organism: (none)
Submitted organism has type: No
Status: INCONCLUSIVE
Confidence: LOW

**Bin 73**

Predicted organism: (none)
Submitted organism has type: No
Status: INCONCLUSIVE
Confidence: LOW
97.504 (34.4	.0)	775708 assembly	1242 Salinimicrobium xinjiangense DSM 19287 (GCA_000423585.1, ASM42358v1)

**Bin 74**

Predicted organism: Tolypothrix tenuis (taxid = 457083, rank = species, lineage = Bacteria; Cyanobacteria; Nostocales; Tolypothrichaceae; Tolypothrix)
Submitted organism has type: No
Status: INCONCLUSIVE
Confidence: LOW
91.847 (45.3	.0) 5101288 assembly	1938 Tolypothrix tenuis PCC 7101 (GCA_002368295.1, ASM236829v1)

**Bin 75**

Predicted organism: (none)
Submitted organism has type: No
Status: INCONCLUSIVE
Confidence: LOW

**Bin 76**

Predicted organism: (none)
Submitted organism has type: No
Status: INCONCLUSIVE
Confidence: LOW

**Bin 77**

Predicted organism: (none)
Submitted organism has type: No
Status: INCONCLUSIVE
Confidence: LOW

**Bin 78**

Predicted organism: (none)
Submitted organism has type: No
Status: INCONCLUSIVE
Confidence: LOW

**Bin 79**

Predicted organism: (none)
Submitted organism has type: No
Status: INCONCLUSIVE
Confidence: LOW

**Bin 80**

Predicted organism: (none)
Submitted organism has type: No
Status: INCONCLUSIVE
Confidence: LOW

**Bin 81**

Predicted organism: (none)
Submitted organism has type: No
Status: INCONCLUSIVE
Confidence: LOW
79.016 (26.2	.0)	528908 assembly	1292 Baaleninema simplex PCC 7105 (GCA_000332355.1, ASM33235v1)

**Bin 82**

Predicted organism: (none)
Submitted organism has type: No
Status: INCONCLUSIVE
Confidence: LOW

**Bin 83**

Predicted organism: (none)
Submitted organism has type: No
Status: INCONCLUSIVE
Confidence: LOW

**Bin 84**

Predicted organism: Nostoc punctiforme (taxid = 272131, rank = species, lineage = Bacteria; Cyanobacteria; Nostocales; Nostocaceae; Nostoc)
Submitted organism has type: No
Status: INCONCLUSIVE
Confidence: LOW
89.714 (52.3	.0)	20828 assembly	1750 Nostoc punctiforme PCC 73102 (GCA_000020025.1, ASM2002v1)

Table S3: Gene models associated with Biofilm formation

| **Description** | **Symbol** | **K** **number** | **Gene** **Models** |
| --- | --- | --- | --- |
| serine 0-acetyltransferase | cysE | K00640 | gnl\|extdb\|tmp_002171,  gnl\|extdb\|tmp_006520,  gnl\|extdb\|tmp_006961,  gnl\|extdb\|tmp_000305,  gnl\|extdb\|tmp_016451,  gnl\|extdb\|tmp_001109,  gnl\|extdb\|tmp_007151,  gnl\|extdb\|tmp_007886,  gnl\|extdb\|tmp_012689,  gnl\|extdb\|tmp_014106,  gnl\|extdb\|tmp_001132,  gnl\|extdb\|tmp_001492,  gnl\|extdb\|tmp_005872,  gnl\|extdb\|tmp_006541,  gnl\|extdb\|tmp_006674,  gnl\|extdb\|tmp_000985,  gnl\|extdb\|tmp_001317,  gnl\|extdb\|tmp_004226,  gnl\|extdb\|tmp_005629,  gnl\|extdb\|tmp_006218 |
| glycogen phosphyrlase | PYG | K00688 | gnl\|extdb\|tmp_009327,  gnl\|extdb\|tmp_006782,  gnl\|extdb\|tmp_002324,  gnl\|extdb\|tmp_004251,  gnl\|extdb\|tmp_004272,  gnl\|extdb\|tmp_005846,  gnl\|extdb\|tmp_006126,  gnl\|extdb\|tmp_007557,  gnl\|extdb\|tmp_010105,  gnl\|extdb\|tmp_000783,  gnl\|extdb\|tmp_014015,  gnl\|extdb\|tmp_001743,  gnl\|extdb\|tmp_011714,  gnl\|extdb\|tmp_013161,  gnl\|extdb\|tmp_014323,  gnl\|extdb\|tmp_015965,  gnl\|extdb\|tmp_017404,  gnl\|extdb\|tmp_017939,  gnl\|extdb\|tmp_003466,  gnl\|extdb\|tmp_003492,  gnl\|extdb\|tmp_004247,  gnl\|extdb\|tmp_004948,  gnl\|extdb\|tmp_006226,  gnl\|extdb\|tmp_006679,  gnl\|extdb\|tmp_007015,  gnl\|extdb\|tmp_007199,  gnl\|extdb\|tmp_010558,  gnl\|extdb\|tmp_011498,  gnl\|extdb\|tmp_012443,  gnl\|extdb\|tmp_004730,  gnl\|extdb\|tmp_008722,  gnl\|extdb\|tmp_009935,  gnl\|extdb\|tmp_009959,  gnl\|extdb\|tmp_013732,  gnl\|extdb\|tmp_014506,  gnl\|extdb\|tmp_000349,  gnl\|extdb\|tmp_001083,  gnl\|extdb\|tmp_001381,  gnl\|extdb\|tmp_002881,  gnl\|extdb\|tmp_004829,  gnl\|extdb\|tmp_005874,  gnl\|extdb\|tmp_007181,  gnl\|extdb\|tmp_007560,  gnl\|extdb\|tmp_001687,  gnl\|extdb\|tmp_001918,  gnl\|extdb\|tmp_007217 |
| cellulose synthase | bcsA | K00694 | gnl\|extdb\|tmp_006796,  gnl\|extdb\|tmp_007011,  gnl\|extdb\|tmp_007319,  gnl\|extdb\|tmp_008422,  gnl\|extdb\|tmp_001067,  gnl\|extdb\|tmp_004810,  gnl\|extdb\|tmp_015071,  gnl\|extdb\|tmp_016632,  gnl\|extdb\|tmp_017793,  gnl\|extdb\|tmp_017826,  gnl\|extdb\|tmp_020100,  gnl\|extdb\|tmp_001230,  gnl\|extdb\|tmp_001873,  gnl\|extdb\|tmp_002162,  gnl\|extdb\|tmp_004616,  gnl\|extdb\|tmp_008840,  gnl\|extdb\|tmp_008986,  gnl\|extdb\|tmp_009002,  gnl\|extdb\|tmp_011350,  gnl\|extdb\|tmp_009513,  gnl\|extdb\|tmp_012386,  gnl\|extdb\|tmp_012521,  gnl\|extdb\|tmp_013055,  gnl\|extdb\|tmp_001770,  gnl\|extdb\|tmp_002131,  gnl\|extdb\|tmp_002577,  gnl\|extdb\|tmp_003296,  gnl\|extdb\|tmp_004283,  gnl\|extdb\|tmp_007547,  gnl\|extdb\|tmp_001160,  gnl\|extdb\|tmp_001201,  gnl\|extdb\|tmp_001615,  gnl\|extdb\|tmp_002198,  gnl\|extdb\|tmp_002513,  gnl\|extdb\|tmp_004738,  gnl\|extdb\|tmp_005559 |
| starch synthase | glgA | K00703 | gnl\|extdb\|tmp_009985,  gnl\|extdb\|tmp_004301,  gnl\|extdb\|tmp_005823,  gnl\|extdb\|tmp_006788,  gnl\|extdb\|tmp_009093,  gnl\|extdb\|tmp_000181,  gnl\|extdb\|tmp_005438,  gnl\|extdb\|tmp_020220,  gnl\|extdb\|tmp_020318,  gnl\|extdb\|tmp_001350,  gnl\|extdb\|tmp_005674,  gnl\|extdb\|tmp_007182,  gnl\|extdb\|tmp_008021,  gnl\|extdb\|tmp_008657,  gnl\|extdb\|tmp_008991,  gnl\|extdb\|tmp_009338,  gnl\|extdb\|tmp_014331,  gnl\|extdb\|tmp_001511,  gnl\|extdb\|tmp_004803,  gnl\|extdb\|tmp_005652,  gnl\|extdb\|tmp_001941,  gnl\|extdb\|tmp_005706 |
| glucose-1-phosphate adenyltransferase | glgC | K00975 | gnl\|extdb\|tmp_000167,  gnl\|extdb\|tmp_003430,  gnl\|extdb\|tmp_005115,  gnl\|extdb\|tmp_000525,  gnl\|extdb\|tmp_004061,  gnl\|extdb\|tmp_013040,  gnl\|extdb\|tmp_001042,  gnl\|extdb\|tmp_002215,  gnl\|extdb\|tmp_004585,  gnl\|extdb\|tmp_005758,  gnl\|extdb\|tmp_009197,  gnl\|extdb\|tmp_008723,  gnl\|extdb\|tmp_015891,  gnl\|extdb\|tmp_015918,  gnl\|extdb\|tmp_001830, gnl\|extdb\|tmp_000836,  gnl\|extdb\|tmp_000304 |
| anthranilate synthase component I | trpE | K01657 | gnl\|extdb\|tmp_002752,  gnl\|extdb\|tmp_004809,  gnl\|extdb\|tmp_010539, gnl\|extdb\|tmp_006191,  gnl\|extdb\|tmp_008273,  gnl\|extdb\|tmp_011663, gnl\|extdb\|tmp_004391,  gnl\|extdb\|tmp_004911 |
| anthranilate synthase component II | TrpG | K01658 | gnl\|extdb\|tmp_014775,  gnl\|extdb\|tmp_001473,  gnl\|extdb\|tmp_004115,  gnl\|extdb\|tmp_014802,  gnl\|extdb\|tmp_015267, gnl\|extdb\|tmp_001155,  gnl\|extdb\|tmp_004413,  gnl\|extdb\|tmp_004631 |
| adenylate cyclase | E4.6.1.1 | K01768 | gnl\|extdb\|tmp_000946,  gnl\|extdb\|tmp_003206,  gnl\|extdb\|tmp_004518,  gnl\|extdb\|tmp_004872,  gnl\|extdb\|tmp_005069,  gnl\|extdb\|tmp_005381,  gnl\|extdb\|tmp_005778,  gnl\|extdb\|tmp_007193,  gnl\|extdb\|tmp_007892,  gnl\|extdb\|tmp_000022,  gnl\|extdb\|tmp_001472,  gnl\|extdb\|tmp_001674,  gnl\|extdb\|tmp_001687,  gnl\|extdb\|tmp_012851,  gnl\|extdb\|tmp_005419,  gnl\|extdb\|tmp_011178,  gnl\|extdb\|tmp_012508,  gnl\|extdb\|tmp_013570,  gnl\|extdb\|tmp_013721,  gnl\|extdb\|tmp_014054,  gnl\|extdb\|tmp_014830,  gnl\|extdb\|tmp_016428,  gnl\|extdb\|tmp_016850,  gnl\|extdb\|tmp_016854,  gnl\|extdb\|tmp_016982,  gnl\|extdb\|tmp_019067,  gnl\|extdb\|tmp_020242,  gnl\|extdb\|tmp_000950,  gnl\|extdb\|tmp_003617,  gnl\|extdb\|tmp_003800,  gnl\|extdb\|tmp_004947,  gnl\|extdb\|tmp_005825,  gnl\|extdb\|tmp_006497,  gnl\|extdb\|tmp_006587,  gnl\|extdb\|tmp_007679,  gnl\|extdb\|tmp_011169,  gnl\|extdb\|tmp_008644,  gnl\|extdb\|tmp_010434,  gnl\|extdb\|tmp_008339,  gnl\|extdb\|tmp_009351,  gnl\|extdb\|tmp_009618,  gnl\|extdb\|tmp_012159,  gnl\|extdb\|tmp_012656,  gnl\|extdb\|tmp_013890,  gnl\|extdb\|tmp_014529,  gnl\|extdb\|tmp_000381,  gnl\|extdb\|tmp_002075,  gnl\|extdb\|tmp_003250,  gnl\|extdb\|tmp_007440,  gnl\|extdb\|tmp_007521,  gnl\|extdb\|tmp_007743,  gnl\|extdb\|tmp_004056,  gnl\|extdb\|tmp_001497,  gnl\|extdb\|tmp_001642,  gnl\|extdb\|tmp_001708,  gnl\|extdb\|tmp_004026,  gnl\|extdb\|tmp_004945,  gnl\|extdb\|tmp_005237,  gnl\|extdb\|tmp_006312,  gnl\|extdb\|tmp_006404 |
| UDP-N-acetylglucosamine 2-epimerase (non-hydrolysing) | wecB | K01791 | gnl\|extdb\|tmp_003714,  gnl\|extdb\|tmp_004402,  gnl\|extdb\|tmp_004429,  gnl\|extdb\|tmp_000764,  gnl\|extdb\|tmp_017475,  gnl\|extdb\|tmp_002658,  gnl\|extdb\|tmp_009134,  gnl\|extdb\|tmp_011473,  gnl\|extdb\|tmp_011676,  gnl\|extdb\|tmp_001910,  gnl\|extdb\|tmp_005981, gnl\|extdb\|tmp_002901,  gnl\|extdb\|tmp_003927 |
| polysaccharide biosynthesis/export protein | wza, gfcE | K01991 | gnl\|extdb\|tmp_002460,  gnl\|extdb\|tmp_003810,  gnl\|extdb\|tmp_005499,  gnl\|extdb\|tmp_006500,  gnl\|extdb\|tmp_006705,  gnl\|extdb\|tmp_009725,  gnl\|extdb\|tmp_010320,  gnl\|extdb\|tmp_010710,  gnl\|extdb\|tmp_013838,  gnl\|extdb\|tmp_013992,  gnl\|extdb\|tmp_016018,  gnl\|extdb\|tmp_004567,  gnl\|extdb\|tmp_008110,  gnl\|extdb\|tmp_004638,  gnl\|extdb\|tmp_006227,  gnl\|extdb\|tmp_009081,  gnl\|extdb\|tmp_009115,  gnl\|extdb\|tmp_009694,  gnl\|extdb\|tmp_012080,  gnl\|extdb\|tmp_012278,  gnl\|extdb\|tmp_006706,  gnl\|extdb\|tmp_002847,  gnl\|extdb\|tmp_001453,  gnl\|extdb\|tmp_003496,  gnl\|extdb\|tmp_007402 |
| general secretion pathway protein E | gspE | K02454 | gnl\|extdb\|tmp_014906,  gnl\|extdb\|tmp_016492 |
| twitching motility two-component system response regulator PilG | pilG | K02657 | gnl\|extdb\|tmp_007932, gnl\|extdb\|tmp_013741,  gnl\|extdb\|tmp_015835,  gnl\|extdb\|tmp_016009, gnl\|extdb\|tmp_009803, gnl\|extdb\|tmp_003522 |
| twitching motility two-component system response regulator PilH | pilH | K02658 | gnl\|extdb\|tmp_005415,  gnl\|extdb\|tmp_004410,  gnl\|extdb\|tmp_000543, gnl\|extdb\|tmp_004021,  gnl\|extdb\|tmp_012834,  gnl\|extdb\|tmp_015943, gnl\|extdb\|tmp_008957, gnl\|extdb\|tmp_003521,  gnl\|extdb\|tmp_005017 |
| twitching motility protein PilJ | pilJ | K02660 | gnl\|extdb\|tmp_009263,  gnl\|extdb\|tmp_004613,  gnl\|extdb\|tmp_005315,  gnl\|extdb\|tmp_005728,  gnl\|extdb\|tmp_009426,  gnl\|extdb\|tmp_014162,  gnl\|extdb\|tmp_016971,  gnl\|extdb\|tmp_018722,  gnl\|extdb\|tmp_000176,  gnl\|extdb\|tmp_001471,  gnl\|extdb\|tmp_009531,  gnl\|extdb\|tmp_009851,  gnl\|extdb\|tmp_012039,  gnl\|extdb\|tmp_013370,  gnl\|extdb\|tmp_016024,  gnl\|extdb\|tmp_001224,  gnl\|extdb\|tmp_008114, gnl\|extdb\|tmp_005242 |
| RNA polymerase nonessential primary-like sigma factor | rpoS | K03087 | gnl\|extdb\|tmp_002592,  gnl\|extdb\|tmp_004166,  gnl\|extdb\|tmp_005407,  gnl\|extdb\|tmp_007449,  gnl\|extdb\|tmp_008508,  gnl\|extdb\|tmp_013572,  gnl\|extdb\|tmp_013744,  gnl\|extdb\|tmp_014411,  gnl\|extdb\|tmp_014812,  gnl\|extdb\|tmp_015210,  gnl\|extdb\|tmp_015241,  gnl\|extdb\|tmp_018950,  gnl\|extdb\|tmp_020049,  gnl\|extdb\|tmp_001691,  gnl\|extdb\|tmp_006156,  gnl\|extdb\|tmp_007840,  gnl\|extdb\|tmp_009192,  gnl\|extdb\|tmp_009987,  gnl\|extdb\|tmp_011492,  gnl\|extdb\|tmp_005944,  gnl\|extdb\|tmp_008316,  gnl\|extdb\|tmp_008656,  gnl\|extdb\|tmp_008812,  gnl\|extdb\|tmp_009274,  gnl\|extdb\|tmp_012160,  gnl\|extdb\|tmp_012644,  gnl\|extdb\|tmp_013748,  gnl\|extdb\|tmp_015733,  gnl\|extdb\|tmp_000935,  gnl\|extdb\|tmp_002050,  gnl\|extdb\|tmp_002660,  gnl\|extdb\|tmp_003478,  gnl\|extdb\|tmp_004726,  gnl\|extdb\|tmp_002926,  gnl\|extdb\|tmp_003888,  gnl\|extdb\|tmp_004346,  gnl\|extdb\|tmp_005250,  gnl\|extdb\|tmp_007089 |
| 3',5'-cyclic-AMP phosphodiesterase | cpdA | K03651 | gnl\|extdb\|tmp_002185,  gnl\|extdb\|tmp_010035, gnl\|extdb\|tmp_012178, gnl\|extdb\|tmp_002509,  gnl\|extdb\|tmp_006981 |
| host factor-I protein | hfq | K03666 | gnl\|extdb\|tmp_010575, gnl\|extdb\|tmp_011943, gnl\|extdb\|tmp_011887 |
| CRP/FNR family transcriptional regulator, cyclic AMP receptor protein | crp | K10914 | gnl\|extdb\|tmp_003429,  gnl\|extdb\|tmp_000315,  gnl\|extdb\|tmp_016471,  gnl\|extdb\|tmp_015138,  gnl\|extdb\|tmp_016528,  gnl\|extdb\|tmp_019890,  gnl\|extdb\|tmp_004686,  gnl\|extdb\|tmp_005541,  gnl\|extdb\|tmp_009009,  gnl\|extdb\|tmp_010224,  gnl\|extdb\|tmp_004571,  gnl\|extdb\|tmp_000683,  gnl\|extdb\|tmp_001050,  gnl\|extdb\|tmp_003171,  gnl\|extdb\|tmp_004741 |
| type VI secretion system protein ImpJ | impJ, vasE | K11893 | gnl\|extdb\|tmp_012125 |
| methyl-accepting chemotaxis protein WspA | wspA | K13487 | gnl\|extdb\|tmp_012518 |
| chemotaxis-related protein WspB | wspB | K13488 | gnl\|extdb\|tmp_014197 |
| two-component system, chemotaxis family, response regulator WspF | wspF | K13491 | gnl\|extdb\|tmp_006170 |
| polysaccharide biosynthesis transport protein | exoP, vspO | K16554 | gnl\|extdb\|tmp_005047,  gnl\|extdb\|tmp_008228,  gnl\|extdb\|tmp_002446,  gnl\|extdb\|tmp_003037,  gnl\|extdb\|tmp_004786,  gnl\|extdb\|tmp_005213,  gnl\|extdb\|tmp_005804,  gnl\|extdb\|tmp_005902,  gnl\|extdb\|tmp_006560,  gnl\|extdb\|tmp_007076,  gnl\|extdb\|tmp_007224,  gnl\|extdb\|tmp_007840,  gnl\|extdb\|tmp_008006,  gnl\|extdb\|tmp_008831,  gnl\|extdb\|tmp_009059,  gnl\|extdb\|tmp_009181,  gnl\|extdb\|tmp_009629,  gnl\|extdb\|tmp_009899,  gnl\|extdb\|tmp_010530,  gnl\|extdb\|tmp_001087,  gnl\|extdb\|tmp_001280,  gnl\|extdb\|tmp_001654,  gnl\|extdb\|tmp_009472,  gnl\|extdb\|tmp_013007,  gnl\|extdb\|tmp_014331,  gnl\|extdb\|tmp_014816,  gnl\|extdb\|tmp_001700,  gnl\|extdb\|tmp_012883,  gnl\|extdb\|tmp_012940,  gnl\|extdb\|tmp_013477,  gnl\|extdb\|tmp_013497,  gnl\|extdb\|tmp_013609,  gnl\|extdb\|tmp_013892,  gnl\|extdb\|tmp_015521,  gnl\|extdb\|tmp_016821,  gnl\|extdb\|tmp_017343,  gnl\|extdb\|tmp_017692,  gnl\|extdb\|tmp_018731,  gnl\|extdb\|tmp_019036,  gnl\|extdb\|tmp_001704,  gnl\|extdb\|tmp_002011,  gnl\|extdb\|tmp_002402,  gnl\|extdb\|tmp_002729,  gnl\|extdb\|tmp_002893,  gnl\|extdb\|tmp_003336,  gnl\|extdb\|tmp_003799,  gnl\|extdb\|tmp_005047,  gnl\|extdb\|tmp_005156,  gnl\|extdb\|tmp_005440,  gnl\|extdb\|tmp_005768,  gnl\|extdb\|tmp_006363,  gnl\|extdb\|tmp_007092,  gnl\|extdb\|tmp_007354,  gnl\|extdb\|tmp_008574,  gnl\|extdb\|tmp_008912,  gnl\|extdb\|tmp_009334,  gnl\|extdb\|tmp_009403,  gnl\|extdb\|tmp_010027,  gnl\|extdb\|tmp_011618,  gnl\|extdb\|tmp_012004,  gnl\|extdb\|tmp_012208,  gnl\|extdb\|tmp_013315,  gnl\|extdb\|tmp_012557,  gnl\|extdb\|tmp_007247,  gnl\|extdb\|tmp_010094,  gnl\|extdb\|tmp_010228,  gnl\|extdb\|tmp_010265,  gnl\|extdb\|tmp_010399,  gnl\|extdb\|tmp_011195,  gnl\|extdb\|tmp_011227,  gnl\|extdb\|tmp_011897,  gnl\|extdb\|tmp_012156,  gnl\|extdb\|tmp_012668,  gnl\|extdb\|tmp_012807,  gnl\|extdb\|tmp_013553,  gnl\|extdb\|tmp_013572,  gnl\|extdb\|tmp_013796,  gnl\|extdb\|tmp_014315,  gnl\|extdb\|tmp_015209,  gnl\|extdb\|tmp_015504,  gnl\|extdb\|tmp_015635,  gnl\|extdb\|tmp_000620,  gnl\|extdb\|tmp_001773,  gnl\|extdb\|tmp_002063,  gnl\|extdb\|tmp_002857,  gnl\|extdb\|tmp_004265,  gnl\|extdb\|tmp_005405,  gnl\|extdb\|tmp_006077,  gnl\|extdb\|tmp_006184,  gnl\|extdb\|tmp_006581,  gnl\|extdb\|tmp_006978,  gnl\|extdb\|tmp_007183,  gnl\|extdb\|tmp_007324,  gnl\|extdb\|tmp_007376,  gnl\|extdb\|tmp_004039,  gnl\|extdb\|tmp_000084,  gnl\|extdb\|tmp_001387,  gnl\|extdb\|tmp_001836,  gnl\|extdb\|tmp_002851,  gnl\|extdb\|tmp_003712,  gnl\|extdb\|tmp_003986,  gnl\|extdb\|tmp_004712,  gnl\|extdb\|tmp_005043,  gnl\|extdb\|tmp_005322,  gnl\|extdb\|tmp_006533,  gnl\|extdb\|tmp_006622 |
| polysaccharide biosynthesis protein PslH | pslH | K21001 | gnl\|extdb\|tmp_008997,  gnl\|extdb\|tmp_003803,  gnl\|extdb\|tmp_007833,  gnl\|extdb\|tmp_015613,  gnl\|extdb\|tmp_016847,  gnl\|extdb\|tmp_000439,  gnl\|extdb\|tmp_002936,  gnl\|extdb\|tmp_004274,  gnl\|extdb\|tmp_006687,  gnl\|extdb\|tmp_015050,  gnl\|extdb\|tmp_015657,  gnl\|extdb\|tmp_006888, gnl\|extdb\|tmp_001173 |

Table S3: Gene models associated with Quorum Sensing

| **Description** | **Symbol** | **K number** | **Gene models** |
| --- | --- | --- | --- |
| trpE anthranilate synthase component I [EC:4.1.3.27] | trpE | K01657 | gnl\|extdb\|tmp_004391, gnl\|extdb\|tmp_004911 |
| glutamate decarboxylase [EC:4.1.1.15] | gadB, gadA, GAD | K01580 | gnl\|extdb\|tmp_011012, gnl\|extdb\|tmp_014064, gnl\|extdb\|tmp_005343 |
| 3-deoxy-7-phosphoheptulonate synthase | aroF, aroG, aroH | K01626 | gnl\|extdb\|tmp_006842, gnl\|extdb\|tmp_007368 |
| anthranilate synthase component I | trpE | K01657 | gnl\|extdb\|tmp_002752, gnl\|extdb\|tmp_004809, gnl\|extdb\|tmp_010539 |
| anthranilate synthase component II | trpG | K01658 | gnl\|extdb\|tmp_014802, gnl\|extdb\|tmp_015267 |
| long-chain acyl-CoA synthetase [EC:6.2.1.3] | ACSL | K01897 | gnl\|extdb\|tmp_002514,  gnl\|extdb\|tmp_003682,  gnl\|extdb\|tmp_004092,  gnl\|extdb\|tmp_004939,  gnl\|extdb\|tmp_005031,  gnl\|extdb\|tmp_005555,  gnl\|extdb\|tmp_005798 |
| branched-chain amino acid transport system ATP-binding protein | livG | K01995 | gnl\|extdb\|tmp_000464,  gnl\|extdb\|tmp_001203 |
| branched-chain amino acid transport system ATP-binding protein | livF | K01996 | gnl\|extdb\|tmp_000182 |
| branched-chain amino acid transport system permease protein | livH | K01997 | gnl\|extdb\|tmp_007038 |
| branched-chain amino acid transport system permease protein | livM | K01998 | gnl\|extdb\|tmp_005973 |
| branched-chain amino acid transport system substrate-binding protein | livK | K01999 | gnl\|extdb\|tmp_006846,  gnl\|extdb\|tmp_007599 |
| peptide/nickel transport system ATP-binding protein | ddpD | K02031 | gnl\|extdb\|tmp_013579,  gnl\|extdb\|tmp_013698,  gnl\|extdb\|tmp_014805,  gnl\|extdb\|tmp_017845,  gnl\|extdb\|tmp_019164,  gnl\|extdb\|tmp_000265,  gnl\|extdb\|tmp_000997,  gnl\|extdb\|tmp_003856,  gnl\|extdb\|tmp_003932,  gnl\|extdb\|tmp_005405,  gnl\|extdb\|tmp_005621,  gnl\|extdb\|tmp_007292,  gnl\|extdb\|tmp_010589,  gnl\|extdb\|tmp_011697,  gnl\|extdb\|tmp_012108 |
| peptide/nickel transport system permease protein | ABC.PE.P | K02033 | gnl\|extdb\|tmp_007329,  gnl\|extdb\|tmp_000363,  gnl\|extdb\|tmp_001354,  gnl\|extdb\|tmp_005117,  gnl\|extdb\|tmp_005324,  gnl\|extdb\|tmp_006172 |
| peptide/nickel transport system permease protein | ABC.PE.P1 | K02034 | gnl\|extdb\|tmp_001293,  gnl\|extdb\|tmp_007011 |
| peptide/nickel transport system substrate-binding protein | ABC.PE.S | K02035 | gnl\|extdb\|tmp_006114 |
| putative spermidine/putrescine transport system substrate-binding protein | ABC.SP.S | K02055 | gnl\|extdb\|tmp_001871 |
| preprotein translocase subunit SecA | secA | K03070 | gnl\|extdb\|tmp_000815,  gnl\|extdb\|tmp_000558,  gnl\|extdb\|tmp_002530,  gnl\|extdb\|tmp_004652,  gnl\|extdb\|tmp_004720,  gnl\|extdb\|tmp_005838 |
| preprotein translocase subunit | secE | K03073 | gnl\|extdb\|tmp_015606,  gnl\|extdb\|tmp_001796 |
| preprotein translocase subunit SecG | SecG | K03075 | gnl\|extdb\|tmp_010610 |
| signal recognition particle subunit SRP54 | SRP54, ffh | K03106 | gnl\|extdb\|tmp_001769,  gnl\|extdb\|tmp_003302 |
| fused signal recognition particle receptor | ftsY | K03110 | gnl\|extdb\|tmp_003815,  gnl\|extdb\|tmp_007378 |
| YidC/Oxa1 family membrane protein insertase | ydC, spoIIIJ, OXA1, ccfA | K03217 | gnl\|extdb\|tmp_004573 |
| host factor-I protein | hfq | K03666 | gnl\|extdb\|tmp_011887 |
| bacterial/archaeal transporter family-2 protein | TC.BAT2 | K09936 | gnl\|extdb\|tmp_001123 |
| CRP/FNR family transcriptional regulator, cyclic AMP receptor protein | crp | K10914 | gnl\|extdb\|tmp_000683,  gnl\|extdb\|tmp_001050,  gnl\|extdb\|tmp_003171,  gnl\|extdb\|tmp_004741 |
| diaminohydroxyphosphoribosylaminopyrimidine deaminase / 5-amino-6-(5-phosphoribosylamino)uracil reductase | ribD | K11752 | gnl\|extdb\|tmp_014874,  gnl\|extdb\|tmp_011070,  gnl\|extdb\|tmp_011580,  gnl\|extdb\|tmp_015275,  gnl\|extdb\|tmp_005285 |
| N-acyl homoserine lactone hydrolase | ahlD, aiiA, attM, blcC | K13075 | gnl\|extdb\|tmp_004381 |
| serine protease | K14645 | K14645 | gnl\|extdb\|tmp_003265,  gnl\|extdb\|tmp_003688 |
| large repetitive protein | bapA | K20276 | gnl\|extdb\|tmp_000394,  gnl\|extdb\|tmp_003043 |
| mannose-binding lectin | bcl | K20527 | gnl\|extdb\|tmp_007324 |
| oligopeptide transport system ATP-binding protein | oppF | K1083 | gln\|xtdb\|tmp_019666 |
| rhamnosyltransferase | rfbF, rhlC | K1290 | MISSING |
| oligopeptide transport system ATP-binding protein | oppD | K15583 |  |
| rhamnosyltransferase subunit B | rhlB | K18101 | MISSING |
| LuxR family transcriptional regulator | raiR | K25873 | gnl\|extdb\|tmp_010628,  gnl\|extdb\|tmp _017967,  gnl\|extdb\|tmp _025873,  gnl\|extdb\|tmp_026581,  gnl\|extdb\|tmp_026585,  gnl\|extdb\|tmp_029011,  gnl\|extdb\|tmp_030422,  gnl\|extdb\|tmp_012176,  gnl\|extdb\|tmp_014828,  gnl\|extdb\|tmp_024923,  gnl\|extdb\|tmp_027010,  gnl\|extdb\|tmp_033322,  gnl\|extdb\|tmp_014501,  gnl\|extdb\|tmp_015089,  gnl\|extdb\|tmp_021317,  gnl\|extdb\|tmp_021500,  gnl\|extdb\|tmp_024808,  gnl\|extdb\|tmp_003105,  gnl\|extdb\|tmp_003107,  gnl\|extdb\|tmp_008059,  gnl\|extdb\|tmp_008089,  gnl\|extdb\|tmp_009914,  gnl\|extdb\|tmp_004548 |

Table S5: Gene models associated with Nitrogen Fixation

| **Description** | **Symbol** | **K number** | **Gene models** |
| --- | --- | --- | --- |
| glutamate dehydrogenase | gdhA | K00261 | gnl\|extdb\|tmp_029999,  gnl\|extdb\|tmp_016328,  gnl\|extdb\|tmp_008838,  gnl\|extdb\|tmp_008839,  gnl\|extdb\|tmp_008840,  gnl\|extdb\|tmp_001969,  gnl\|extdb\|tmp_006204 |
| glutamate dehydrogenase | gdhA | K00262 | gnl\|extdb\|tmp_010647 |
| glutamate synthase | gltB | K00265 | gnl\|extdb\|tmp_004992,  gnl\|extdb\|tmp_006971 |
| glutamate synthase | gltD | K00266 | gnl\|extdb\|tmp_001253 |
| glutamate synthase | gltS, GLU | K00284 | gnl\|extdb\|tmp_018018,  gnl\|extdb\|tmp_018019,  gnl\|extdb\|tmp_018020,  gnl\|extdb\|tmp_018608,  gnl\|extdb\|tmp_018609,  gnl\|extdb\|tmp_013163,  gnl\|extdb\|tmp_013164,  gnl\|extdb\|tmp_013165,  gnl\|extdb\|tmp_019211,  gnl\|extdb\|tmp_019212,  gnl\|extdb\|tmp_019214 |
| ferredoxin nitrite reductase | nirA | K00366 | gnl\|extdb\|tmp_027560,  gnl\|extdb\|tmp_030554,  gnl\|extdb\|tmp_022688,  gnl\|extdb\|tmp_022689,  gnl\|extdb\|tmp_025332,  gnl\|extdb\|tmp_018903,  gnl\|extdb\|tmp_020224,  gnl\|extdb\|tmp_022867,  gnl\|extdb\|tmp_022868,  gnl\|extdb\|tmp_003323 |
| ferredoxin nitrate reductase | narB | K00367 | gnl\|extdb\|tmp_010257,  gnl\|extdb\|tmp_022691,  gnl\|extdb\|tmp_020213,  gnl\|extdb\|tmp_020214,  gnl\|extdb\|tmp_008516,  gnl\|extdb\|tmp_008517,  gnl\|extdb\|tmp_000053,  gnl\|extdb\|tmp_000054 |
| nitrate reductase | narG, narZ, nxrA | K00370 | gnl\|extdb\|tmp_005404 |
| nitronate monooxygenase | ncdw, npd | K00459 | gnl\|extdb\|tmp_016776,  gnl\|extdb\|tmp_004936 |
| formamidase | E3.5.1.49 | K01455 | gnl\|extdb\|tmp_010727,  gnl\|extdb\|tmp_029679 |
| nitrilase | E3.5.5.1 | K01501 | gnl\|extdb\|tmp_033069 |
| carbonic anhydrase | can, cynT | K01673 | gnl\|extdb\|tmp_010778,  gnl\|extdb\|tmp_032344,  gnl\|extdb\|tmp_012875,  gnl\|extdb\|tmp_026756,  gnl\|extdb\|tmp_026757,  gnl\|extdb\|tmp_030311,  gnl\|extdb\|tmp_012693,  gnl\|extdb\|tmp_015116,  gnl\|extdb\|tmp_016118,  gnl\|extdb\|tmp_016132,  gnl\|extdb\|tmp_018541,  gnl\|extdb\|tmp_018542,  gnl\|extdb\|tmp_021919,  gnl\|extdb\|tmp_021918,  gnl\|extdb\|tmp_004340,  gnl\|extdb\|tmp_006758,  gnl\|extdb\|tmp_006759,  gnl\|extdb\|tmp_007738,  gnl\|extdb\|tmp_000797,  gnl\|extdb\|tmp_000949,  gnl\|extdb\|tmp_005126 |
| carbonic anhydrase | cah | K01674 | gnl\|extdb\|tmp_011074,  gnl\|extdb\|tmp_011073 |
| cyanate lyase | cynS | K01725 | gnl\|extdb\|tmp_033075,  gnl\|extdb\|tmp_019537,  gnl\|extdb\|tmp_005658 |
| glutamine synthethase | glnA, GLUL | K01915 | gnl\|extdb\|tmp_030941,  gnl\|extdb\|tmp_032377,  gnl\|extdb\|tmp_008339,  gnl\|extdb\|tmp_009237,  gnl\|extdb\|tmp_003759,  gnl\|extdb\|tmp_004361 |
| nitrate/nitrite transporter | NRT, narK, nrtP, nasA | K02575 | gnl\|extdb\|tmp_010256,  gnl\|extdb\|tmp_022690,  gnl\|extdb\|tmp_033264,  gnl\|extdb\|tmp_033265,  gnl\|extdb\|tmp_020216,  gnl\|extdb\|tmp_008621,  gnl\|extdb\|tmp_009321 |
| nitrogenase molybdenum-iron protein alpha chain | nifD | K02586 | gnl\|extdb\|tmp_028508,  gnl\|extdb\|tmp_013104,  gnl\|extdb\|tmp_007646,  gnl\|extdb\|tmp_008032,  gnl\|extdb\|tmp_008033,  gnl\|extdb\|tmp_008205,  gnl\|extdb\|tmp_003343 |
| nitrogenase iron protein | nifH | K02588 | gnl\|extdb\|tmp_026743,  gnl\|extdb\|tmp_026744,  gnl\|extdb\|tmp_028298,  gnl\|extdb\|tmp_021957,  gnl\|extdb\|tmp_002720,  gnl\|extdb\|tmp_007645,  gnl\|extdb\|tmp_008030,  gnl\|extdb\|tmp_000194,  gnl\|extdb\|tmp_006568 |
| nitrogenase molybdenum-iron protein beta chain | nifK | K02591 | gnl\|extdb\|tmp_028509,  gnl\|extdb\|tmp_002477,  gnl\|extdb\|tmp_008774,  gnl\|extdb\|tmp_001711 |
| nitrate/nitrite transport system substrate binding protein | nrtA | K15576 | gnl\|extdb\|tmp_011105,  gnl\|extdb\|tmp_033072,  gnl\|extdb\|tmp_028577,  gnl\|extdb\|tmp_012536,  gnl\|extdb\|tmp_012537,  gnl\|extdb\|tmp_014469,  gnl\|extdb\|tmp_014470,  gnl\|extdb\|tmp_020007,  gnl\|extdb\|tmp_020222,  gnl\|extdb\|tmp_020223,  gnl\|extdb\|tmp_001045 |
| nitrate/nitrite transport system permease protein | nrtB, nasE, cynB | K15577 | gnl\|extdb\|tmp_011106,  gnl\|extdb\|tmp_033073,  gnl\|extdb\|tmp_028578,  gnl\|extdb\|tmp_012058,  gnl\|extdb\|tmp_012525,  gnl\|extdb\|tmp_012538,  gnl\|extdb\|tmp_012539,  gnl\|extdb\|tmp_014467,  gnl\|extdb\|tmp_020220,  gnl\|extdb\|tmp_020221,  gnl\|extdb\|tmp_001044 |
| nitrate/nitrite transport system ATP binding protein | nrtC, nasD | K15578 | gnl\|extdb\|tmp_011107,  gnl\|extdb\|tmp_033074,  gnl\|extdb\|tmp_028576,  gnl\|extdb\|tmp_012535,  gnl\|extdb\|tmp_014468,  gnl\|extdb\|tmp_020008,  gnl\|extdb\|tmp_020009,  gnl\|extdb\|tmp_020219,  gnl\|extdb\|tmp_001043 |
| nitrate/nitrite transport system ATP binding protein | nrtD, cynD | K15579 | gnl\|extdb\|tmp_020217,  gnl\|extdb\|tmp_020218,  gnl\|extdb\|tmp_009319,  gnl\|extdb\|tmp_009320 |
| Nonheterocysts |  |  |  |
| Nitrogen fixation protein | nifX | K02596 | gnl\|extdb:tmp_028514, gnl\|extdb:tmp_001800, gnl\|extdb:tmp_015269 |
| Associated nitrogen fixation protein | nifX-associated nitrogen fixation protein |  | gnl\|extdb:tmp_028515, gnl\|extdb:tmp_001714, gnl\|extdb:tmp_015271, gnl\|extdb:tmp_015270 |
| Nitrogenase cofactor biosynthesis protein | nifB | K02585 | gnl\|extdb:tmp_008837, gnl\|extdb:tmp_026728, gnl\|extdb:tmp_026734 |
| nitrogen fixation protein NifZ | nifZ | K02594 | gnl\|extdb:tmp_012197, gnl\|extdb:tmp_027418, gnl\|extdb:tmp_005710, gnl\|extdb:tmp_007663 |
| nitrogen fixation protein NifT | nifT | K02593 | gnl\|extdb:tmp_012198, gnl\|extdb:tmp_027419, gnl\|extdb:tmp_005709 |
| nitrogenase molybdenum-cofactor synthesis protein NifE | nifEN | K02587 | gnl\|extdb:tmp_015266, gnl\|extdb:tmp_001710 |
| nitrogenase molybdenum-iron protein NifN | nifN | K02592 | gnl\|extdb:tmp_015268, gnl\|extdb:tmp_004277, gnl\|extdb:tmp_003560 |
| NifU family protein | nifU | K04488 | gnl\|extdb:tmp_026724, gnl\|extdb:tmp_021955 |
| Nirogenase iron-molybdenum cofactor synthesis protein | nifE | K02587 | gnl\|extdb:tmp_028511 |
| nitrogenase molybdenum-iron protein NifN | nifN | K02592 | gnl\|extdb:tmp_028513, gnl\|extdb:tmp_003560 |
| homocitrate synthase NifV [EC:2.3.3.14] | nifV | K02594 | gnl\|tmp_012196, gnl\|tmp_027417 |
| cysteine desulfurase [EC:2.8.1.7] | nifS | K04487 | gnl\|extdb:tmp_026737 |
| urease subunit alpha | UreA | K01428 | gnl\|extdb\|tmp_029711,  gnl\|extdb\|tmp_014298 |
| urease accessory protein | UreD | K03190 | gnl\|extdb\|tmp_030959,  gnl\|extdb\|tmp_019833,  gnl\|extdb\|tmp_014295,  gnl\|extdb\|tmp_005741 |
| urease accessory protein | UreE | K03187 | gnl\|extdb\|tmp_025770,  gnl\|extdb\|tmp_022574 |
| urease accessory protein | UreF | K03188 | gnl\|extdb\|tmp_025771,  gnl\|extdb\|tmp_022573 |
| urease acessory protein | UreG | K03189 | gnl\|extdb\|tmp_030960,  gnl\|extdb\|tmp_025772,  gnl\|extdb\|tmp_014294,  gnl\|extdb\|tmp_022572 |
| urease subunit beta | UreB | K01429 | gnl\|extdb\|tmp_030961,  gnl\|extdb\|tmp_014293 |
| urease subunit gamma | UreC | No K# assigned | gnl\|extdb\|tmp_030962,  gnl\|extdb\|tmp_019834,  gnl\|extdb\|tmp_014292,  gnl\|extdb\|tmp_005742 |
| ammonification of amino acids |  |  |  |
| asparaginase | ASRGL-1 | K13051 | gnl\|extdb\|tmp_021385,  gnl\|extdb\|tmp_021386,  gnl\|extdb\|tmp_003548,  gnl\|extdb\|tmp_029170,  gnl\|extdb\|tmp_031006,  gnl\|extdb\|tmp_019863,  gnl\|extdb\|tmp_027905,  gnl\|extdb\|tmp_027906,  gnl\|extdb\|tmp_027909,  gnl\|extdb\|tmp_027910,  gnl\|extdb\|tmp_017411,  gnl\|extdb\|tmp_017769,  gnl\|extdb\|tmp_021129 |
| glutaminase | TGMX | K05622 | gnl\|extdb\|tmp_023778,  gnl\|extdb\|tmp_010254,  gnl\|extdb\|tmp_011036,  gnl\|extdb\|tmp_011160,  gnl\|extdb\|tmp_029326,  gnl\|extdb\|tmp_029494,  gnl\|extdb\|tmp_022111,  gnl\|extdb\|tmp_027137,  gnl\|extdb\|tmp_029921,  gnl\|extdb\|tmp_012329,  gnl\|extdb\|tmp_016962,  gnl\|extdb\|tmp_017117,  gnl\|extdb\|tmp_017118,  gnl\|extdb\|tmp_018253,  gnl\|extdb\|tmp_018808,  gnl\|extdb\|tmp_019543,  gnl\|extdb\|tmp_021285, |

Table S6: Gene models associated with the sequestering and/or transport of metals

| **Description** | **Symbol** | **K number** | **Gene models** |
| --- | --- | --- | --- |
| PhoD-like phosphatase | PhoD | K01113 | gnl\|extdb\|tmp_026528,  gnl\|extdb\|tmp_013876,  gnl\|extdb\|tmp_016745,  gnl\|extdb\|tmp_020129,  gnl\|extdb\|tmp_006427,  gnl\|extdb\|tmp_007217,  gnl\|extdb\|tmp_009935 |
| Alkaline phosphatase D | PhoD | K01113 | gnl\|extdb\|tmp_007060,  gnl\|extdb\|tmp_000741,  gnl\|extdb\|tmp_002818,  gnl\|extdb\|tmp_031076,  gnl\|extdb\|tmp_016746,  gnl\|extdb\|tmp_016748,  gnl\|extdb\|tmp_020130,  gnl\|extdb\|tmp_021644,  gnl\|extdb\|tmp_021645,  gnl\|extdb\|tmp_021646  gnl\|extdb\|tmp_007059 |
| Metal transporter | CNNM | K16302 | gnl\|extdb\|tmp_011921 |
| Nramp family divalent metal transporter | SMF | K12346 | gnl\|extdb\|tmp_011888,  gnl\|extdb\|tmp_005137 |
| Zip family metal transporter | TC.ZIP,  ZupT,ZRT3, ZIP2, ZIP | K07238 | gnl\|extdb\|tmp_003522,  gnl\|extdb\|tmp_026424,  gnl\|extdb\|tmp_004434 |
| TonB siderophore receptor | TC.FEV.OM | K02014 | gnl\|extdb\|tmp_002010,  gnl\|extdb\|tmp_011049,  gnl\|extdb\|tmp_026381,  gnl\|extdb\|tmp_029066,  gnl\|extdb\|tmp_030640,  gnl\|extdb\|tmp_013299,  gnl\|extdb\|tmp_024362,  gnl\|extdb\|tmp_025306,  gnl\|extdb\|tmp_028440,  gnl\|extdb\|tmp_030153,  gnl\|extdb\|tmp_020995,  gnl\|extdb\|tmp_022762,  gnl\|extdb\|tmp_004957,  gnl\|extdb\|tmp_008746 |
| Siderphore transporter | SIT1 | No K# assigned | gnl\|extdb\|tmp_006418,  gnl\|extdb\|tmp_026381 |
| Siderphore ABC transporter | YfiYZ/YfhA/YusV | No K# assigned | gnl\|extdb\|tmp_005492,  gnl\|extdb\|tmp_011048,  gnl\|extdb\|tmp_026382,  gnl\|extdb\|tmp_013301,  gnl\|extdb\|tmp_024260,  gnl\|extdb\|tmp_024341,  gnl\|extdb\|tmp_025316,  gnl\|extdb\|tmp_028441,  gnl\|extdb\|tmp_013261,  gnl\|extdb\|tmp_020695,  gnl\|extdb\|tmp_020991,  gnl\|extdb\|tmp_022751,  gnl\|extdb\|tmp_023475,  gnl\|extdb\|tmp_001633,  gnl\|extdb\|tmp_001635,  gnl\|extdb\|tmp_004416,  gnl\|extdb\|tmp_008059,  gnl\|extdb\|tmp_000606 |
| Multicopper oxidase family protein | MCO | No K# assigned | gnl\|extdb\|tmp_003877 |
| Multicopper oxidase domain containing protein |  | No K# assigned | gnl\|extdb\|tmp_002015,  gnl\|extdb\|tmp_003877,  gnl\|extdb\|tmp_004009,  gnl\|extdb\|tmp_004783,  gnl\|extdb\|tmp_004784,  gnl\|extdb\|tmp_007285,  gnl\|extdb\|tmp_008132,  gnl\|extdb\|tmp_000404,  gnl\|extdb\|tmp_004075,  gnl\|extdb\|tmp_014059,  gnl\|extdb\|tmp_011915,  gnl\|extdb\|tmp_011916,  gnl\|extdb\|tmp_011919,  gnl\|extdb\|tmp_011920,  gnl\|extdb\|tmp_013214,  gnl\|extdb\|tmp_013215,  gnl\|extdb\|tmp_015045,  gnl\|extdb\|tmp_015047 |

Supplementary Table S7: Genes associated with Pigments

| **Description** | **Symbol** | **K number** | **Gene models** |
| --- | --- | --- | --- |
| Scytonemin biosynthesis cyclase decarboxylase | ScyC | No K# assigned | gnl\|extdb\|tmp_007802  gnl\|extdb\|tmp_025745  gnl\|extdb\|tmp_007902 |
| Scytonemin biosynthesis cyclase decarboxylase | ScyA | No K# assigned | gnl\|extdb\|tmp_025743 |
| Scytonemin biosynthesis cyclase decarboxylase | ScyF | No K# assigned | gnl\|extdb\|tmp_025748 |
| UbiA-like protein EboC | EcoC | No K# assigned | gnl\|extdb\|tmp_025751 |
| Carotenoid oxidase | CrtO | No K# assigned | gnl\|extdb\|tmp_013790 |
| Orange carotenoid protein | Ocp | No K# assigned | gnl\|extdb\|tmp_014572  gnl\|extdb\|tmp_018150  gnl\|extdb\|tmp_020564  gnl\|extdb\|tmp_020580  gnl\|extdb\|tmp_001461  gnl\|extdb\|tmp_006160  gnl\|extdb\|tmp_007245  gnl\|extdb\|tmp_031997  gnl\|extdb\|tmp_014697  gnl\|extdb\|tmp_027358  gnl\|extdb\|tmp_032508 |
| 1-deoxy-D-xylulose-5-phosphate reductoisomerase activity | Dxr | K00099 | gnl\|extdb\|tmp_013173  gnl\|extdb\|tmp_029386 |
| phytoene desaturase | Crtl, PDS | K02293 | gnl\|extdb\|tmp_030876 |
| Phytoene synthase | crtB | K02291 | gnl\|extdb\|tmp_015072  gnl\|extdb\|tmp_015073 |
| Carotene isomerase | crtH | K09835 | gnl\|extdb\|tmp_027155  gnl\|extdb\|tmp_029748 |
| geranylgeranyl diphosphate reductase | GGR | K17830 |  |
| 9,9'-di-cis-zeta-carotene desaturase | ZDS | K00514 | gnl\|extdb\|tmp_019825  gnl\|extdb\|tmp_025403  gnl\|extdb\|tmp_021812  gnl\|extdb\|tmp_010315 |
| Phycocyanin alpha chain | cpcA | K02284 | gnl\|extdb\|tmp_030485  gnl\|extdb\|tmp_030491  gnl\|extdb\|tmp_032865  gnl\|extdb\|tmp_032873  gnl\|extdb\|tmp_032874  gnl\|extdb\|tmp_016703  gnl\|extdb\|tmp_021309  gnl\|extdb\|tmp_007569 |
| Phycocyanin beta chain | cpcB | K02285 | gnl\|extdb\|tmp_030492  gnl\|extdb\|tmp_033104  gnl\|extdb\|tmp_015843  gnl\|extdb\|tmp_031575  gnl\|extdb\|tmp_032864  gnl\|extdb\|tmp_014911  gnl\|extdb\|tmp_016702  gnl\|extdb\|tmp_021309  gnl\|extdb\|tmp_024754  gnl\|extdb\|tmp_007595 |
| Allophycocyanin alpha subunit | apcA | K02092 | gnl\|extdb\|tmp_029304  gnl\|extdb\|tmp_030887  gnl\|extdb\|tmp_033105  gnl\|extdb\|tmp_022564 |
|  |  |  | gnl\|extdb\|tmp_021237 |
| Allophycocyanin subunit beta | apcB | K02093 | gnl\|extdb\|tmp_030888  gnl\|extdb\|tmp_030889  gnl\|extdb\|tmp_030942  gnl\|extdb\|tmp_021239  gnl\|extdb\|tmp_003756 |
| Allophycocyanin subunit alpha-B | apcD | K02095 | gnl\|extdb\|tmp_031922  gnl\|extdb\|tmp_026633  gnl\|extdb\|tmp_025163  gnl\|extdb\|tmp_004558 |
| Chromophore lyase CpcT/CpeT | cpcT/CpeT | No K# assigned | gnl\|extdb\|tmp_012210  gnl\|extdb\|tmp_026966  gnl\|extdb\|tmp_028583  gnl\|extdb\|tmp_032938  gnl\|extdb\|tmp_011973  gnl\|extdb\|tmp_013499  gnl\|extdb\|tmp_013506  gnl\|extdb\|tmp_017016  gnl\|extdb\|tmp_020787  gnl\|extdb\|tmp_006266  gnl\|extdb\|tmp_028830  gnl\|extdb\|tmp_023294  gnl\|extdb\|tmp_006492  gnl\|extdb\|tmp_029598 |

Table S8: Gene models associated with desiccation tolerance.

| Description | Symbol | K Number | Gene Models |
| --- | --- | --- | --- |
| Sugars |  |  |  |
| sedoheptulose 7-phosphate cyclase | slig | K19969 | gnl\|extdb:tmp_030397, gnl\|extdb:tmp_032658, gnl\|extdb:tmp_018838 |
| malto-oligosyltrehalose synthase | treY | K06044 | gnl\|extdb\|tmp_010631,  gnl\|extdb\|tmp_016529,  gnl\|extdb\|tmp_025438,  gnl\|extdb\|tmp_025439,  gnl\|extdb\|tmp_017184 |
| malto-oligosyltrehalose trehalohydrolase | treZ | K01236 | gnl\|extdb\|tmp_010632,  gnl\|extdb\|tmp_016530 |
| trehalase family glycosidase | Not Assigned | Not Assigned | gnl\|extdb:tmp_010636, gnl\|extdb:tmp_010637, gnl\|extdb:tmp_016527, gnl\|extdb:tmp_016528, gnl\|extdb:tmp_025437, gnl\|extdb:tmp_017186 |
| sucrose-phosphate phosphatase | Not Assigned | Not Assigned | gnl\|extdb\|tmp_025230,  gnl\|extdb\|tmp_018283,  gnl\|extdb\|tmp_018721,  gnl\|extdb\|tmp_003702,  gnl\|extdb\|tmp_026359 |
| sucrose synthase activity | Not Found | Not Found | gnl\|extdb\|tmp_018723 |
| Extrapolysaccharides Biosynthesis |  |  |  |
| serine O-acetyltransferase | cysE | K00640 | gnl\|extdb\|tmp_012192,  gnl\|extdb\|tmp_018086,  gnl\|extdb\|tmp_023465,  gnl\|extdb\|tmp_027414,  gnl\|extdb\|tmp_032444,  gnl\|extdb\|tmp_012381,  gnl\|extdb\|tmp_017069,  gnl\|extdb\|tmp_017070,  gnl\|extdb\|tmp_004988,  gnl\|extdb\|tmp_005140,  gnl\|extdb\|tmp_008636,  gnl\|extdb\|tmp_003395,  gnl\|extdb\|tmp_006367 |
| UDP-GlcNAc:undecaprenyl-phosphate/decaprenyl-phosphate GlcNAc-1-phosphate transferase | wecA, tagO, rfe | K02851 | gnl\|extdb\|tmp_033000,  gnl\|extdb\|tmp_032697,  gnl\|extdb\|tmp_015498,  gnl\|extdb\|tmp_019355,  gnl\|extdb\|tmp_023949,  gnl\|extdb\|tmp_003386,  gnl\|extdb\|tmp_002014 |
| putative colanic acid biosynthesis glycosyltransferase | wcaI | K03208 | gnl\|extdb\|tmp_027659,  gnl\|extdb\|tmp_012222,  gnl\|extdb\|tmp_025752,  gnl\|extdb\|tmp_015803,  gnl\|extdb\|tmp_021126,  gnl\|extdb\|tmp_005821,  gnl\|extdb\|tmp_007813,  gnl\|extdb\|tmp_007814 |
| putative colanic acid biosynthesis acetyltransferase | wcaF | K03818 | gnl\|extdb\|tmp_030636,  gnl\|extdb\|tmp_018845,  gnl\|extdb\|tmp_009066,  gnl\|extdb\|tmp_010050,  gnl\|extdb\|tmp_002575 |
| succinoglycan biosynthesis protein | exoO | K16555 | gnl\|extdb\|tmp_018097,  gnl\|extdb\|tmp_022535 |
| succinoglycan biosynthesis protein | exoA | K16557 | gnl\|extdb\|tmp_030044 |
| colanic acid/amylovoran/stewartan biosynthesis glycosyltransferase | wcaL, amsK, cpsK | K16703 | gnl\|extdb\|tmp_025975,  gnl\|extdb\|tmp_031860,  gnl\|extdb\|tmp_031872,  gnl\|extdb\|tmp_031875,  gnl\|extdb\|tmp_012837,  gnl\|extdb\|tmp_012840,  gnl\|extdb\|tmp_018081,  gnl\|extdb\|tmp_018083,  gnl\|extdb\|tmp_019334,  gnl\|extdb\|tmp_019335,  gnl\|extdb\|tmp_028305,  gnl\|extdb\|tmp_030232 |
| colanic acid/amylovoran biosynthesis protein | wcaK, amsJ | K16710 | gnl\|extdb\|tmp_012835,  gnl\|extdb\|tmp_012836 |
| mannuronan synthase | alg8 | K19290 | gnl\|extdb\|tmp_012149 |
| alginate O-acetyltransferase complex protein AlgI | algI | K19294 | gnl\|extdb\|tmp_027724,  gnl\|extdb\|tmp_029045,  gnl\|extdb\|tmp_015731,  gnl\|extdb\|tmp_021972,  gnl\|extdb\|tmp_022938,  gnl\|extdb\|tmp_022939,  gnl\|extdb\|tmp_008542,  gnl\|extdb\|tmp_008543 |
| polysaccharide biosynthesis protein PslH | pslH | K21001 | gnl\|extdb\|tmp_028388,  gnl\|extdb\|tmp_015510,  gnl\|extdb\|tmp_015514,  gnl\|extdb\|tmp_021846,  gnl\|extdb\|tmp_001737,  gnl\|extdb\|tmp_004153,  gnl\|extdb\|tmp_004154 |
| TEMP |  |  |  |
| chaperone DnaK | dnaK | K04043 | gnl\|extdb:tmp_017237, gnl\|extdb:tmp_027981, gnl\|extdb:tmp_033490, gnl\|extdb:tmp_014975, gnl\|extdb:tmp_026301, gnl\|extdb:tmp_027598, gnl\|extdb:tmp_029360, gnl\|extdb:tmp_017237, |
| chaperone DnaJ | dnaJ | K03686 | gnl\|extdb:tmp_020321, gnl\|extdb:tmp_008193, gnl\|extdb:tmp_026300, gnl\|extdb:tmp_032840 |
| Chaperonim GroEL | groEL | K04077 | gnl\|extdb:tmp_001184, gnl\|extdb:tmp_005723, gnl\|extdb:tmp_006885, gnl\|extdb:tmp_030624, gnl\|extdb:tmp_030945, gnl\|extdb:tmp_019830, gnl\|extdb:tmp_032696, gnl\|extdb:tmp_019357, gnl\|extdb:tmp_003385, gnl\|extdb:tmp_005507, gnl\|extdb:tmp_007809,  gnl\|tmp_009584 |
| Hsp70 family protein | HSP70 | K14017 | gnl\|extdb:tmp_004240, gnl\|extdb:tmp_004242, gnl\|extdb:tmp_029133, gnl\|extdb:tmp_030731, gnl\|extdb:tmp_033489, gnl\|extdb:tmp_020323, gnl\|extdb:tmp_023056, gnl\|extdb:tmp_000498, gnl\|extdb:tmp_004653, gnl\|extdb:tmp_004737, gnl\|extdb:tmp_007518, gnl\|extdb:tmp_008370, gnl\|extdb:tmp_008371, gnl\|extdb:tmp_020323, gnl\|extdb:tmp_023056, gnl\|extdb:tmp_023058, gnl\|extdb:tmp_000498, gnl\|extdb:tmp_004653, gnl\|extdb:tmp_004737, gnl\|extdb:tmp_007518, gnl\|extdb:tmp_008370, gnl\|extdb:tmp_008370, gnl\|extdb:tmp_008371, gnl\|extdb:tmp_008372, gnl\|extdb:tmp_008373, gnl\|extdb:tmp_009076, gnl\|extdb:tmp_009485 |
| aquaporins | aqpZ | K06188 | gnl\|extdb:tmp_000219, gnl\|extdb:tmp_000584, gnl\|extdb:tmp_027546, gnl\|extdb:tmp_030814,  gnl\|extdb:tmp_013251 |
| Antioxidant enzyme systems |  |  |  |
| superoxide dismutase | sodN | K00518 | gnl\|extdb:tmp_018912, gnl\|extdb:tmp_024895, gnl\|extdb:tmp_001957, gnl\|extdb:tmp_002450, gnl\|extdb:tmp_003139, gnl\|extdb:tmp_007846, gnl\|extdb:tmp_010133, gnl\|extdb:tmp_010398, gnl\|extdb:tmp_030828, gnl\|extdb:tmp_011801, gnl\|extdb:tmp_013761, gnl\|extdb:tmp_018255 |
| glutathione peroxidase | gpx | K00432 | gnl\|extdb:tmp_019721, gnl\|tmp_019725, gnl\|extdb:tmp_032148, gnl\|extdb:tmp_023467, gnl\|extdb:tmp_032859 |
| manganese catalase family protein | Ini | K07217 | gnl\|extdb:tmp_019946, gnl\|extdb:tmp_025116, gnl\|extdb:tmp_002415, gnl\|extdb:tmp_000405, gnl\|extdb:tmp_010324, gnl\|extdb:tmp_031986, gnl\|extdb:tmp_014113, gnl\|extdb:tmp_016494, gnl\|extdb:tmp_025797, gnl\|extdb:tmp_014365, gnl\|extdb:tmp_018215 |
| spermidine synthase (polyamine synthesis) | speE | K00797 | gnl\|extdb:tmp_010488, gnl\|extdb:tmp_011148, gnl\|extdb:tmp_011149, gnl\|extdb:tmp_023179, gnl\|extdb:tmp_030598, |
| Akinete formation |  |  |  |
| heterocyst differentiation master regulator | hetR | Not Assigned | gnl\|extdb:tmp_024561, gnl\|extdb:tmp_007444, gnl\|extdb:tmp_000712 |
| HetP family heterocyst commitment protein | hetP | Not Assigned | gnl\|extdb:tmp_007495, gnl\|extdb:tmp_027838, gnl\|extdb:tmp_028989, gnl\|extdb:tmp_018629, gnl\|extdb:tmp_026777, gnl\|extdb:tmp_011469, gnl\|extdb:tmp_015453, gnl\|extdb:tmp_020031, gnl\|extdb:tmp_024561, gnl\|extdb:tmp_002620, gnl\|extdb:tmp_007444 |
